# Photosynthesis-derived carbon gates cell cycle activation to enable hormone-autonomous shoot regeneration

**DOI:** 10.64898/2026.06.25.734677

**Authors:** Yetkin Çaka Ince, Arika Takebeyashi, Akira Iwase, Johanna Krahmer, Aurore Chetelat, Mitsuhiro Aida, Lieven De Veylder, Keiko Sugimoto

**Affiliations:** RIKEN Center for Sustainable Resource Science, Yokohama 230-0045, Japan; University of Copenhagen, Department for Plant and Environmental Sciences, 1871 Frederiksberg-C, Denmark; Department of Plant Molecular Biology, University of Lausanne, CH-1015 Lausanne, Switzerland; Faculty of Advanced Science and Technology (FAST), Kumamoto University, 2-39-1 Kurokami, Chuo-ku, Kumamoto, Japan; International Research Center for Agricultural and Environmental Biology, Kumamoto University, 2-39-1 Kurokami, Chuo-ku, Kumamoto, Japan; Department of Plant Biotechnology and Bioinformatics, Ghent University, 9052 Ghent, Belgium; VIB Center for Plant Systems Biology, 9052 Ghent, Belgium; Department of Biological Sciences, Graduate School of Science, The University of Tokyo, Bunkyo-ku, Tokyo, 113-0033 Japan

## Abstract

Shoot regeneration is a powerful model for cell fate reprogramming but how it occurs in nature remains poorly understood because studies in *Arabidopsis thaliana* conventionally rely on *in vitro* assays supplemented with exogenous hormones and sugars. In this study, we established the Hormone-autonomY Direct Regeneration Assay (HYDRA) in which removal of the shoot apical meristem (SAM) initiates shoot regeneration from the cotyledon-hypocotyl boundary domain without hormone or sugar supplementation. We show that photosynthesis-derived carbon and the boundary domain are two separable but convergent requirements for shoot regeneration in HYDRA. Carbon availability increases in the boundary domain where it activates cell cycle progression via the RETINOBLASTOMA-RELATED1 (RBR1) pathway. Carbon deprivation blocks regeneration despite induction of SAM marker genes, indicating that carbon-dependent cell cycle activation is a limiting factor for regeneration. In parallel, perturbation of the boundary domain or its regulators reduces regeneration despite sufficient carbon, indicating that boundary domain identity is independently required. Additionally, exogenous carbon supply overcomes the requirement for SAM removal to induce shoot formation, indicating that carbon availability also acts as an initiation cue. Together, this study reveals an inherent capacity for hormone-autonomous shoot regeneration and identifies photosynthesis-derived carbon as a central regulator of this process.

## INTRODUCTION

Plants possess a remarkable capacity to regenerate organs and even entire individuals from differentiated cells. This ability not only provides a framework for studying cell fate reprogramming but also underpins a wide range of applications, including genetic transformation across diverse species and clonal propagation of elite genotypes. Shoot regeneration is particularly critical, as it enables the development of reproductive organs and recovery of whole plants. Although some species can autonomously regenerate shoots, *Arabidopsis thaliana* is considered to lack this capacity and requires exogenous hormone supplementation for efficient shoot regeneration (Amano et al., 2020; Amutha et al., 2009; Damodaran and Strader, 2024; Filipović et al., 2015; Lee et al., 2004; Okazaki et al., 2025; Perica and Berljak, 1996; Pozueta-Romero et al., 2001; Rivers and Marcotrigiano, 1994; Tanimoto and Harada, 1982). Consequently, hormone-induced assays have become the standard approach to study shoot regeneration (Ikeuchi et al., 2019; Ikeuchi et al., 2016). While these assays have advanced our molecular understanding of shoot regeneration, the mechanisms underlying hormone-autonomous shoot regeneration remain poorly defined.

Hormone-induced assays typically use a two-step culture system (Valvekens et al., 1988). First, wounded explants are incubated on auxin-rich callus-inducing medium (CIM) to promote production of pluripotent callus. Subsequently, the explants are transferred to cytokinin-rich shoot-inducing medium (SIM) to regenerate new shoots. Both of these media also include exogenously supplied sugars (Valvekens et al., 1988). Cell proliferation is an essential step for any regeneration process (Che et al., 2007; Fan et al., 2012; Kareem et al., 2015; Lee et al., 2025; Zhai and Xu, 2021). Auxin, cytokinin, and sugars all function in the activation of cell proliferation partly through the CYCLIN D (CYCD) - CYCLIN-DEPENDENT KINASE (CDK)- RETINOBLASTOMA-RELATED1 (RBR1) - E2 promoter-binding factor (E2F) pathway (hereafter referred to as the RBR1 pathway) (Dewitte et al., 2007; Garza-Aguilar et al., 2017; Harashima et al., 2013; Hirano et al., 2008; Liu et al., 2018; Magyar et al., 2005; Perilli et al., 2013; Riou-Khamlichi et al., 1999; Riou-Khamlichi et al., 2000). In this pathway, CYCDs, particularly CYCD3;1, activate CDKs, leading to phosphorylation and inactivation of RBR1 (Magyar et al., 2012; Menges et al., 2006; Nowack et al., 2012). KIP-RELATED PROTEINs (KRPs) inhibit CDKs, maintaining RBR1 in a hypophosphorylated and active state (Magyar et al., 2012; Verkest et al., 2005; Zhao et al., 2017). In its active state, RBR1 binds E2F transcription factors and represses genes associated with cell cycle progression (De Veylder et al., 2002; Francis, 2007). Thus, RBR1 inactivation promotes activation of cell proliferation programs.

Auxin and cytokinin also have distinct roles in regeneration (Ikeuchi et al., 2019; Ikeuchi et al., 2016; Ikeuchi et al., 2013). Auxin promotes the acquisition of shoot competence mainly from xylem pericycle-like cells via genetic programs associated with lateral root primordium and root meristem development (Che et al., 2007; Fan et al., 2012; Kareem et al., 2015; Rosspopoff et al., 2017; Sugimoto et al., 2010; Zhai and Xu, 2021). Following the acquisition of competence, cytokinin drives the conversion to shoot fate by inducing the shoot apical meristem (SAM) regulator *WUSCHEL* (*WUS*) and establishes a *de novo* shoot meristem on SIM (Zhang et al., 2017).

Wounding provides an additional regulatory input and the *WOUND INDUCED DEDIFFERENTIATION 1* (*WIND1*)-mediated pathway plays major roles in this control. *WIND1* is required for wound-induced cell fate conversion and subsequently contributes to shoot regeneration (Iwase et al., 2017; Iwase et al., 2015; Iwase et al., 2011). It also induces *ENHANCER OF SHOOT REGENERATION 1* (*ESR1*), which together with *ESR2* promotes SAM establishment and activates key meristem regulators, including *WUS*, *SHOOT MERISTEMLESS* (*STM*), *CUP SHAPED COTYLEDON 1* (*CUC1*), and *CUC2* (Ikeda et al., 2006; Ikeda et al., 2021; Iwase et al., 2017).

Importantly, STM and CUCs are meristem regulators closely linked to shoot competence, particularly during the initiation of axillary meristems (AMs) from boundary domains (Aida and Tasaka, 2006; Balkunde et al., 2017; Guo et al., 2025; Hepworth and Pautot, 2015; Keller et al., 2006; Long and Barton, 2000; Shi et al., 2016; Zhang et al., 2018). Boundary domains are narrow regions between the SAM and emerging leaves or cotyledons, or between adjacent shoot organs such as leaves and stems. They often show restricted growth and reduced cell division, yet retain competence for AM initiation (Aida and Tasaka, 2006; Hepworth and Pautot, 2015). Genetic studies have demonstrated that *STM, LATERAL SUPPRESSOR (LAS), BREVIPEDICELLUS-LIKE (BLR)*, and *KNOTTED1-LIKE HOMEOBOX GENE 6 (KNAT6)* act redundantly in boundary domain formation between cotyledons and hypocotyl. *CUCs* operate upstream of these regulators (Aida et al., 2020). *STM* emerges as a key component in this network, as the expression of an immobilized form of *STM* is sufficient to induce ectopic leaf initiation from the cotyledon boundary (Balkunde et al., 2017). Additionally, *STM* marks shoot progenitor identity in hormone-induced regeneration assays (Varapparambath et al., 2022; Zhai et al., 2026), highlighting its conserved role in conferring shoot competence across multiple contexts.

Recently, light has emerged as a key environmental cue that influences shoot regeneration in hormone-induced assays (Chen et al., 2024; Dai et al., 2022; Ince and Sugimoto, 2023). Light is perceived by photoreceptors, including phytochromes (PHY; red/far-red light), cryptochromes (CRY; blue light), and UVR8 (UV-B light) (Galvao and Fankhauser, 2015; Legris et al., 2019). The CONSTITUTIVE PHOTOMORPHOGENIC 1 (COP1) acts downstream of these photoreceptors as a central repressor by targeting the key photomorphogenesis transcription factor ELONGATED HYPOCOTYL 5 (HY5) for degradation (Galvao and Fankhauser, 2015). Mutations in *phyA*, *phyB*, *CRY1*, and *HY5* reduce shoot regeneration, whereas *cop1* mutants show enhanced regeneration, particularly under limited light (Chen et al., 2024; Li et al., 2025; Nameth et al., 2013). The effects of light signaling are mediated, at least in part, through interactions with hormone pathways (Cluis et al., 2004; Li et al., 2025; Nameth et al., 2013). This is consistent with the established role of light in regulating auxin and cytokinin responses across diverse developmental contexts (Chory et al., 1994; de Wit et al., 2016; Vandenbussche et al., 2007; Yoshida et al., 2018; Zdarska et al., 2015). However, this tight coupling makes it difficult to fully assess the contribution of light to shoot regeneration in the presence of exogenous hormones.

Exogenous carbon resources in the hormone-induced shoot regeneration assays introduce an additional limitation, since they make it difficult to assess the contribution of endogenous carbon supply. Because external sugars can substitute for photosynthesis-derived carbon, their presence may mask the contribution of light through photosynthesis and prevent direct testing of sugar requirements without a no-sugar control. The role of carbon is clearer in root regeneration systems, which operate without exogenous hormones or sugars (Omary et al., 2023; Sena et al., 2009). Root tip regeneration depends on sugars transported from photosynthetic organs (Matosevich et al., 2026). Light is also necessary to provide carbon resources in *de novo* root regeneration from leaf explants (Chen et al., 2014). Additionally, sugars regulate lateral root initiation, where light acts mainly through photosynthesis rather than signaling (Kircher and Schopfer, 2023). Consistently, we previously showed that photosynthesis remains limiting for shoot regeneration despite intact light signaling, even under sugar supplementation (Chen et al., 2024). Additionally, the importance of photosynthesis-related gene expression is documented in tomato shoot regeneration (Larriba et al., 2021). These observations suggest that the photosynthetic contribution of light to shoot regeneration may be underestimated, highlighting the need for a shoot regeneration assay in *Arabidopsis thaliana* that operates without exogenous inputs.

In this study, we establish a new regeneration assay in *Arabidopsis thaliana* in which shoots regenerate efficiently without exogenous hormones or sugars. In this assay, we show that SAM removal acts as the key initiation signal and triggers an increase in carbon availability in the boundary domain between the cotyledon and hypocotyl. Increased carbon availability promotes cell cycle progression through the RBR1 pathway, with light supporting this process primarily by supplying carbon through photosynthesis. The meristem regulator *STM* is pre-expressed in this domain and is required for shoot regeneration together with other boundary-associated regulators such as *LAS*. We further demonstrate that exogenous carbon supply can bypass the requirement for SAM removal to initiate shoot regeneration. Together, these findings reveal that *Arabidopsis thaliana* possesses an inherent capacity for shoot regeneration, in which carbon availability acts as an early determinant through cell cycle activation.

## RESULTS

### *Arabidopsis thaliana* can regenerate shoots in the absence of exogenous hormones

To develop an assay for hormone-autonomous shoot regeneration in *Arabidopsis thaliana*, we reasoned that the pre-existing shoot apical meristem (SAM) must be completely removed so that newly regenerated shoot meristems can be readily detected. We also aimed to preserve seedling integrity as much as possible to allow other physiological processes to proceed. We therefore removed the entire shoot apex, including the SAM, together with one cotyledon, while preserving the connection between the remaining cotyledon and hypocotyl (Fig. 1A,B). Immediately after dissection, *CLAVATA3* (*CLV3*) reporter signal (Prunet et al., 2017) was restricted to the removed shoot apex, confirming successful SAM removal from the remaining explant (Fig. 1C). We therefore refer to the removed shoot apex as the “SAM explant.” In the remaining explant, after two days (2d) culture on hormone-free Murashige and Skoog (MS) medium post-dissection, both *CLV3* and *WUS* reporter (Tucker et al., 2008) expression became detectable at the cotyledon–hypocotyl boundary, indicating *de novo* shoot meristem formation (Fig. 1D). Continued culture without exogenous hormones, even only on water-soaked filter paper, resulted in consistent formation of multiple shoots, visible 3–7 days post-dissection (Fig. 1E,F; Supplementary Fig. 1A). These observations illustrate that *Arabidopsis thaliana* seedlings possess hormonal autonomy for shoot regeneration. We therefore named this system Hormone-autonomY Direct Regeneration Assay (HYDRA), inspired by the formation of multiple shoots, and hereafter refer to the remaining explant as the “HYDRA explant.”

**Figure 1.**
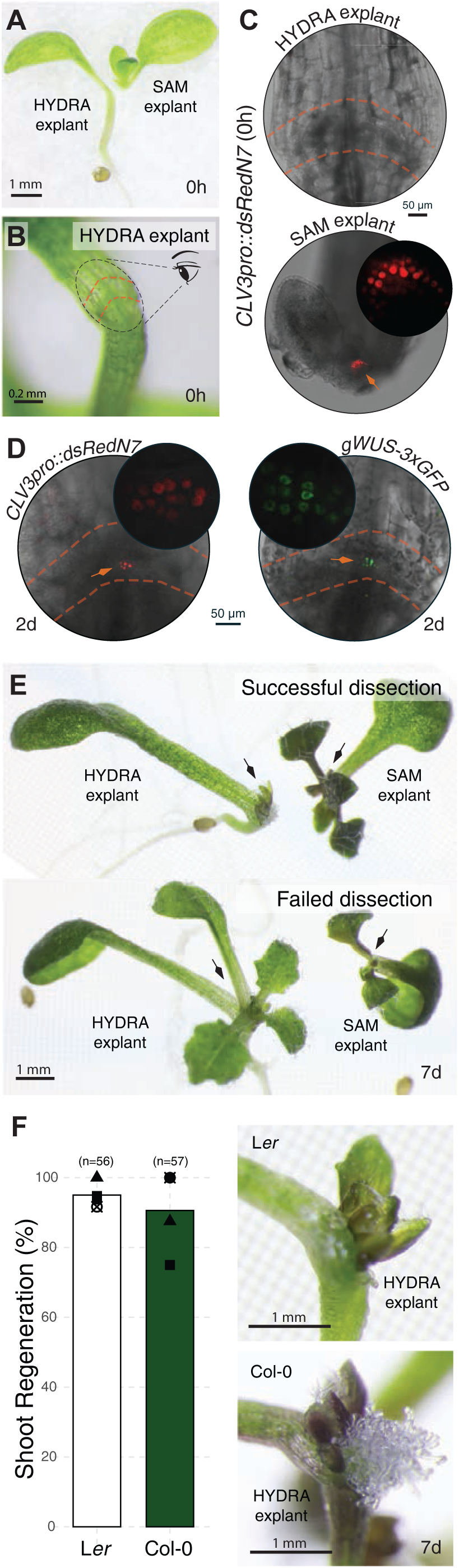
*Arabidopsis thaliana* regenerates *de novo* shoots from the cotyledon-hypocotyl boundary without exogenous hormones. (A) Representative image showing HYDRA dissection in a 7-day-old light-grown Arabidopsis seedling. (B) Close-up of the HYDRA explant immediately after dissection. The region between dashed orange line marks the cotyledon (top) - hypocotyl (bottom) boundary and corresponds to the regions shown in microscopy panels at (C-E). (C) Expression of SAM reporter *CLV3pro::dsRedN7* in HYDRA (top) and SAM (bottom) explants 0h after dissection. Images are maximum projections (top) and single optical sections (bottom). (D) Expression of SAM reporters *CLV3pro::dsRedN7* and *gWUS-3xGFP* in the HYDRA explant 2 days (2d) after dissection. Images are single optical sections. (E) Examples of successful (top) and failed (bottom) dissection experiments. Arrows on SAM explants mark continued (top) or ceased (bottom) leaf initiation. Arrow heads on HYDRA explants mark delayed (i.e. regenerated) (top) or immediate (bottom) leaf initiation. (F) Quantification (left) and representative images (right) for regenerated HYDRA explants 7d after dissection in L*er* and Col-0. Regeneration is scored as presence or absence of shoots. Each dot with different shapes corresponds to a biological replicate. Bars show average of these values. Total sample size (n) is indicated. Seedlings were dissected at 7d after germination and cultured on hormone-free MS medium under continuous light.

To exclude the possibility that the regenerated shoots originated from residual SAM cells due to incomplete dissection, we co-cultured the SAM explant alongside each seedling as a dissection control (Fig. 1E). Successful dissection was indicated by continued leaf development and secondary leaf initiation in the SAM explant, while regenerating shoots appeared in HYDRA explants only 3–7 days post-dissection. In contrast, incomplete removal of the entire SAM was characterized by cessation of growth in the SAM explant, whereas residual SAM cells in the HYDRA explant immediately initiated growth of a pair of leaves that became visible within 1–2 days. These leaves were larger than *de novo* HYDRA shoots, allowing us to identify such cases as “failed dissections” and exclude them from further analysis (Fig. 1E).

We tested HYDRA for the commonly used L*er* and Col-0 ecotypes, both of which showed regeneration frequencies above 90%, indicating that shoot regeneration in HYDRA is a robust event rather than a rare phenomenon (Fig. 1F). Notably, shoot regeneration in HYDRA was also observed in other *Brassicaceae* species, such as *Brassica rapa* and *Brassica napus,* as well as several non*-Brassicaceae* species (Supplementary Figs. 1B, C), suggesting that this regeneration response is conserved across a broad range of dicots.

To establish a consistent quantification strategy, we first considered the morphology and timing of regenerated shoot development. While multiple shoots often regenerated and developed into morphologically normal plants, in some cases one shoot became dominant while others remained dormant (Supplementary Fig. 1D). We therefore decided to quantify shoot regeneration when shoots were clearly visible but before secondary leaf emergence (Fig. 1F). However, regenerated shoots frequently did not display the canonical two-leaf pattern. Enlarged or fused meristems often initiated more than two leaves, and multiple meristems were frequently present within a confined area. As counting individual “shoots” was difficult and prone to interpretation, we instead quantified shoot regeneration based on the number of regenerated leaves (Supplementary Fig. 1E). Only outermost leaves at early initiation stages were counted at 7 days post-dissection, minimizing the influence of subsequent outgrowth.

To test whether this approach reflects differences in meristem size and number, we used the *clv3-2* mutant, in which loss of *CLV3* enlarges the SAM (Clark et al., 1995). In this mutant, the presence of multiple shoots and increased leaf production from enlarged and often fused meristems was pronounced (Supplementary Fig. 1F). Consistently, *clv3-2* showed a significant increase in regenerated leaf number compared to the WT background, supporting the validity of our leaf-based quantification strategy.

### Shoot regeneration in HYDRA tolerates substantial perturbations in endogenous auxin and cytokinin

Auxin and cytokinin are well-established regulators of shoot regeneration in hormone-induced assays (Ikeuchi et al., 2019; Schneider et al., 2019; Smeringai et al., 2023). Given that shoot regeneration in HYDRA occurs without the supply of exogenous hormones, the contribution of endogenous hormone pathways is of particular interest. To assess the role of auxin, we analyzed mutants and chemical inhibitors targeting the major indole-3-acetic acid (IAA) biosynthesis pathway, in which TRYPTOPHAN AMINOTRANSFERASE OF ARABIDOPSIS/TAA1- RELATED (TAA1/TAR) enzymes catalyze the rate-limiting step (Zhao, 2014). The *tar1-1 tar2-1* mutant (Stepanova et al., 2008) did not show reduced shoot regeneration in HYDRA under mock treatment (Supplementary Fig. 2A). Similarly, treatment with the TAA1/TAR inhibitor L-kynurenine (KYN) did not affect regeneration in the wild-type background. However, combining KYN with *tar1-1 tar2-1* reduced regeneration in a dose-dependent manner, although substantial regeneration still occurred (Supplementary Fig. 2A). In addition, treatment with the auxin transport inhibitor N-1-naphthylphthalamic acid (NPA) reduced regeneration only at high concentrations (Supplementary Fig. 2A). Together, these results indicate that auxin contributes to shoot regeneration in HYDRA. However, the persistence of regeneration despite substantial perturbation of auxin biosynthesis and transport suggests that this contribution is buffered by redundancy among pathway components.

We next examined the cytokinin receptor double mutant *ahk3-7 cre1-2,* affecting *ARABIDOPSIS HISTIDINE KINASE 3 (AHK3)* and *CYTOKININ RESPONSE 1 (CRE1)*, which is unable to regenerate shoots in hormone-induced regeneration assays (Pernisova et al., 2018; Riefler et al., 2006). However, this mutant did not exhibit reduced regeneration in HYDRA, suggesting redundancy within cytokinin signaling components (Supplementary Fig. 2B). We therefore used an inducible *CYTOKININ OXIDASE7* (*35S::XVE>>CKX7,* i.e. *CKX7i)* line, in which increased expression of *CKX7* promotes cytokinin degradation and thereby broadly reduces cytokinin signaling (Ye et al., 2021). Induction of *CKX7i* reduced shoot regeneration compared to mock treatment, indicating that cytokinin contributes to the process (Supplementary Fig. 2B). Notably, substantial regeneration persisted, demonstrating tolerance to reduced cytokinin signaling. Together, these results indicate that the endogenous auxin and cytokinin pathways contribute to shoot regeneration in HYDRA, but that the system is relatively tolerant to perturbations in these pathways.

### Shoot regeneration is initiated by SAM removal and requires the boundary domain

We observed that wound-induced callus (WIC) formation accompanies shoot regeneration in HYDRA, particularly in Col-0 but not in L*er* (Fig. 1F). To assess the contribution of WIC in HYDRA, we used the dominant negative *WIND1-SRDX* line (Iwase et al., 2011), which exhibited reduced WIC formation (Supplementary Fig. 3A). Despite this reduction, *WIND1-SRDX* regenerated shoots similarly to WT, indicating that WIC formation is largely uncoupled from shoot regeneration in HYDRA.

Shoot regeneration in HYDRA and *WUS/CLV3* reporter expression were spatially restricted to the cotyledon–hypocotyl boundary domain (Fig. 1D, F). Boundary domains are regions known to harbor cells that are competent for axillary meristem formation (Aida and Tasaka, 2006; Hepworth and Pautot, 2015). Although no established meristems are present in *A. thaliana* cotyledon boundary domain (Grbic and Bleecker, 2000), we hypothesized that the capacity to form new meristems may nevertheless be retained. To test this hypothesis, we performed serial dissections to generate explants with or without the boundary domain and assessed their regeneration capacity (Fig. 2A-C). Cotyledon and hypocotyl explants containing the boundary domain showed regeneration comparable to HYDRA explants, whereas removal of the boundary domain completely abolished shoot regeneration (Figs. 2B, C). In contrast, explants containing only the boundary domain regenerated shoots efficiently (Figs. 2B, C), indicating that the boundary domain is essential for shoot regeneration in HYDRA.

**Figure 2.**
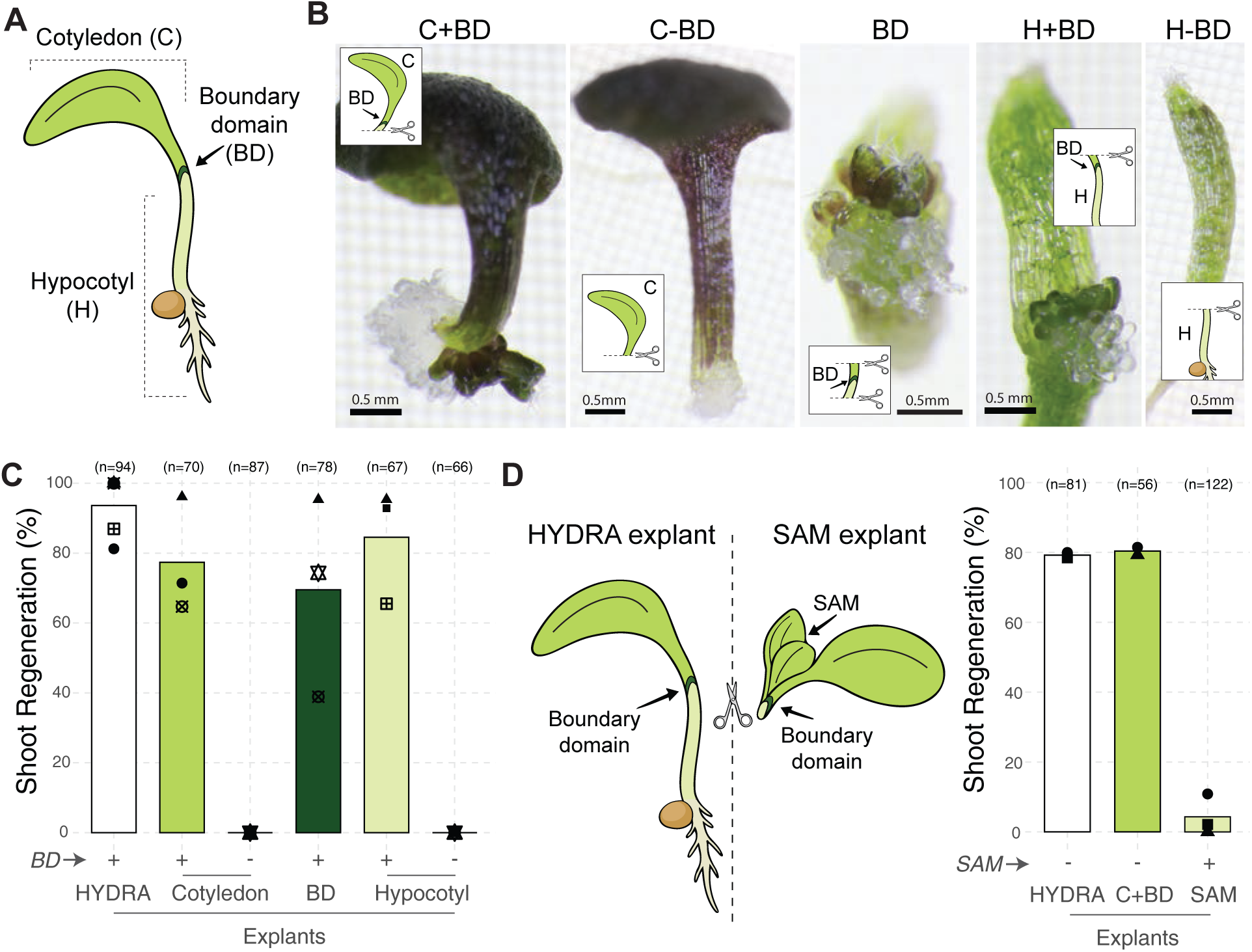
Shoot regeneration is initiated by SAM removal and requires the boundary domain. (A) Schematic of HYDRA explant with marked structures. (B) Representative images of explants with (+BD) or without boundary domain (-BD). (C) The impact of *boundary domain (BD)* presence on shoot regeneration in explants. (D) Schematic of dissections (left) and the impact of *shoot apical meristem (SAM)* presence on shoot regeneration in explants (right). Media were supplemented with sucrose in (B, C) but without sucrose in (D). Regeneration is scored as presence or absence of shoots. Each dot with different shapes corresponds to a biological replicate. Bars show average of these values. Total sample size (n) is indicated.

To further test whether exogenous cytokinin could uncover latent shoot regeneration capacity in WIC in the absence of the boundary domain, we transferred WIC formed from cotyledon explants lacking the boundary domain onto SIM (Supplementary Fig. 3B). However, no shoot regeneration was observed, consistent with our previous findings on WIC derived from hypocotyl explants (Hung et al., 2026). Taken together, these results indicate that shoot regeneration capacity in HYDRA is confined to the boundary domain. This raised the question of whether SAM explants, which also retain the boundary domain and experience similar wounding cues, might also be capable of regenerating shoots (Fig. 2D). To test this, we compared regeneration efficiency between HYDRA and SAM explants, which differ primarily in the presence of the SAM. Unlike HYDRA explants, SAM explants failed to regenerate shoots (Fig. 2D). Given that cotyledon explants containing the boundary domain, but lacking the hypocotyl and root, regenerated comparably to HYDRA explants (Fig. 2C, D), these results indicate that the difference in regeneration capacity between HYDRA and SAM explants is due to the SAM presence. This suggests that the SAM imposes apical dominance that suppresses regeneration from the boundary domain, and that removal of this dominance acts as a cue initiating shoot regeneration in HYDRA.

### Carbon resources are required for shoot regeneration in HYDRA

Light has been shown to be an important determinant of shoot regeneration in hormone-induced assays (Chen et al., 2024; Dai et al., 2022; Ince and Sugimoto, 2023). We therefore tested its role in HYDRA during the regeneration phase. Shoot regeneration was entirely inhibited in darkness (Fig. 3A), indicating that light is essential for shoot regeneration in HYDRA. To distinguish whether this requirement reflects light signaling or photosynthesis-derived carbon, we examined the effects of light quality and quantity. Red, blue, and white light of equal intensity supported similar regeneration, whereas increasing light intensity enhanced regeneration (Fig. 3B). Additionally, a reduced red to far-red (R/FR) ratio, that mimics vegetative shade conditions (Fiorucci and Fankhauser, 2017), did not affect shoot regeneration (Supplementary Fig. 4A,B). These results indicate that light quantity, but not quality, is important for shoot regeneration in HYDRA.

**Figure 3.**
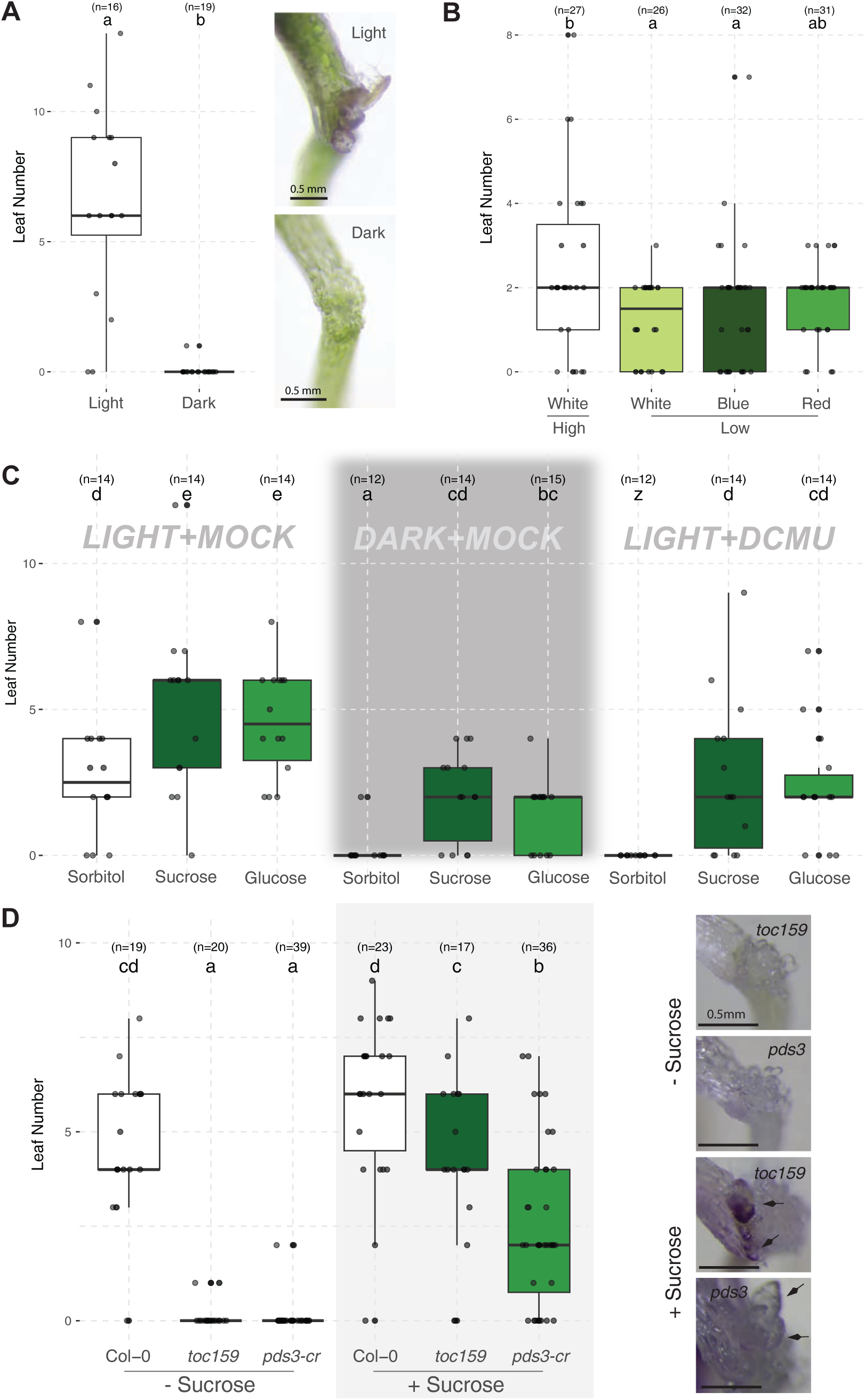
Carbon resources are required for shoot regeneration in HYDRA. (A) Shoot regeneration in continuous light or dark. (B) Shoot regeneration in high (40 µmol/m²s) or low (10 µmol/m²s) light. (C) Shoot regeneration in light or dark with or without DCMU (10 µM) and sugars. (D) Shoot regeneration in photosynthesis mutants with or without sucrose. Treatments were applied during the regeneration phase. Individual data points are shown as dots. Boxplots represent the distribution of the data, the center line indicates the median, the box corresponds to the interquartile range (25th to 75th percentile), and whiskers extend to 1.5 × the interquartile range. Different letters indicate statistical significance (P<0.05), “z” indicates all values are 0 and excluded from statistics. Sample size (n) is indicated.

The dependence on light intensity suggested a dominant role for photosynthetic carbon supply. To test this, we treated seedlings with DCMU (3-(3,4-dichlorophenyl)-1,1-dimethylurea) which inhibits photosynthesis. DCMU completely blocked shoot regeneration under light, phenocopying the dark treatment (Fig. 3C). Supplying metabolizable sucrose or glucose restored regeneration under both dark and DCMU conditions, whereas the osmotic control sorbitol did not (Fig. 3C). Consistent with these results, genetic impairment of photosynthesis also inhibited shoot regeneration. The photosynthesis mutants *toc159* (Kubis et al., 2004) and *pds3-cr* were impaired in shoot regeneration, a phenotype that was rescued by exogenous sucrose (Fig. 3D; Supplementary Fig. 4F). Together, these results demonstrate that photosynthesis-derived carbon availability underlies the requirement for light in HYDRA.

To assess the contribution of light signaling, we analyzed mutants in major light signaling pathways. Consistent with the light quality results, the photoreceptor mutants *phyB-9-BC*, *phyA-211 phyB-9-BC* (Reed et al., 1994; Yoshida et al., 2018), and *cry1-304 cry2-1* (Mockler et al., 1999) regenerated similarly to WT (Supplementary Fig. 4C). In the signaling pathway downstream of photoreceptors, both *cop1* (Ang and Deng, 1994) and its target *hy5-2* (Lee et al., 2011) also showed normal regeneration (Supplementary Fig. 4D, E). To further test whether light signaling can promote regeneration independently of carbon availability, we examined *cop1* mutants in darkness supplemented with sucrose. Under these conditions, where the carbon supply is equal and light signaling is constitutively active in *cop1*, regeneration remained similar to WT (Supplementary Fig. 4E). These results indicate that the activation of major light signaling pathways does not measurably contribute to shoot regeneration in HYDRA. Together, these findings establish carbon availability as a major determinant of shoot regeneration in HYDRA, with light acting primarily through photosynthesis rather than through canonical light signaling pathways.

### Transcriptomics link carbon availability to cell cycle progression in HYDRA

To identify transcriptional events associated with carbon availability during shoot regeneration in HYDRA, we performed RNA sequencing (RNA-seq) on HYDRA explants treated with either mock or DCMU (Fig. 4A). Samples were collected at 6 and 30 hours (h) post-dissection at the same zeitgeber time to capture the early transcriptional events preceding visible *WUS* and *CLV3* reporter expression (Fig. 1D). To enrich for transcriptional signals associated with the shoot meristem formation domain, we collected tissue specifically from the cotyledon–hypocotyl boundary region after removing the surrounding cotyledon and hypocotyl tissues. As a negative control, uncut seedlings were dissected in the same way but immediately frozen. Principal component analysis (PCA) showed tight clustering of biological replicates, confirming reproducibility among biological replicates (Supplementary Fig. 5A). At 6h, mock- and DCMU-treated samples clustered closely in PCA space, indicating minimal early effects of DCMU at this time point. In contrast, at the 30h timepoint, the mock and DCMU samples were clearly separated, revealing a pronounced effect of carbon availability. We therefore focused on the 30h time point to assess carbon-dependent responses in HYDRA.

**Figure 4.**
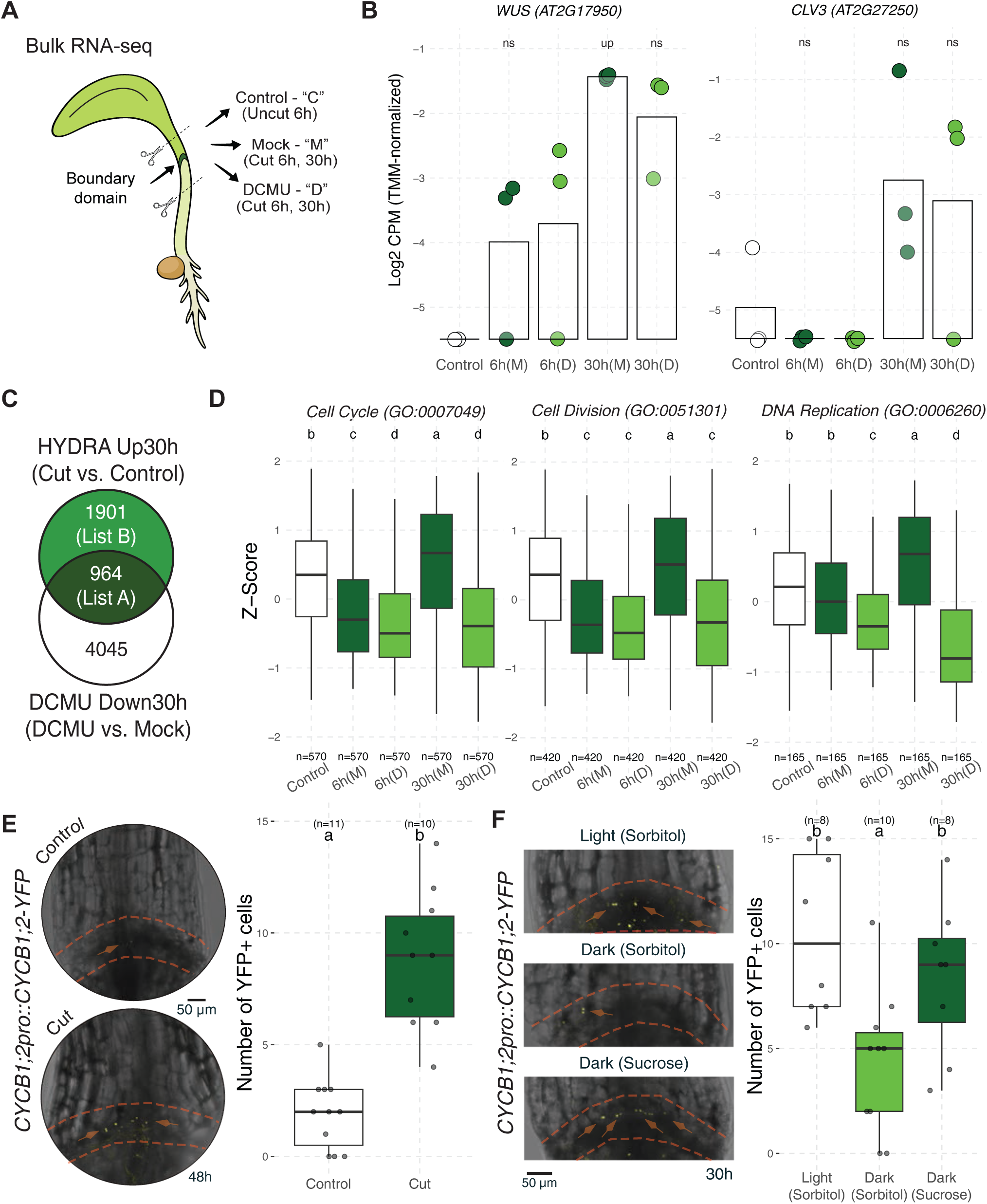
Transcriptomics link carbon supply to cell cycle progression in HYDRA. (A) Experimental setup for transcriptome analysis in HYDRA. (B) Expression of SAM genes *WUS* and *CLV3*. ns: not significant, up: upregulated (FDR <0.05). (C) Venn diagram comparing HYDRA-up and DCMU-downregulated genes (30h) (FC >1, FDR <0.05). List A: HYDRA up & DCMU down regulated genes (carbon-dependent), List B: HYDRA specifically upregulated genes (carbon-independent). (D) Z-scores of all genes that are expressed in our dataset for given GO terms. Gene number in each term expressed in HYDRA dataset is indicated as “n”. (E) Representative images and quantification of CYCB1;2-YFP signal in cut or control (uncut) samples (48h). (F) CYCB1;2-YFP signal in cut samples with indicated treatments (30h). (E, F) Images are maximum projections of Z-stacks. The area between orange line marks the boundary domain. Treatments were applied during the regeneration phase. Boxplots represent the distribution of the data, the center line indicates the median, the box corresponds to the interquartile range (25th to 75th percentile), and whiskers extend to 1.5 × the interquartile range. Different letters indicate statistical significance (P < 0.05). Sample size (n) is indicated.

To determine whether carbon limitation directly affects shoot meristem establishment itself, we first examined the expression of the key shoot meristem markers *WUS* and *CLV3*. *WUS* expression was detectable already at 6h and increased strongly by 30h, whereas *CLV3* expression was detectable only at 30h (Fig. 4B). Notably, despite the complete inhibition of shoot regeneration by DCMU, neither *WUS* nor *CLV3* expression was suppressed (Fig. 4B). This strongly suggests that carbon deprivation does not block the initial transcriptional activation of shoot meristem markers, suggesting that carbon availability acts either downstream of or in parallel to meristem initiation.

To identify the mechanisms by which carbon deprivation limits shoot regeneration, we compared genes upregulated in HYDRA and downregulated by DCMU at 30h (fold change — FC >1, FDR <0.05; Supplementary Table 1). This analysis identified 964 genes that are induced in HYDRA in a carbon-dependent manner (List A, Fig. 4C). Gene Ontology (GO) analysis of these genes revealed strong enrichment for cell cycle, cell division, and related processes (Supplementary Table 2). Importantly, this pattern was not restricted to significantly regulated genes. When considering all genes annotated to the cell cycle, cell division, and DNA replication GO terms, including upregulated, downregulated, and unchanged genes, their overall expression distribution showed a coordinated peak at 30h in mock-treated HYDRA samples and broad suppression by DCMU (Fig. 4D). Notably, the DCMU-dependent downregulation trend was also present at 6h. These results indicate that carbon availability promotes transcriptional activation of cell cycle genes in HYDRA.

To assess whether this transcriptional response is enriched in the boundary domain, we compared our dataset with a set of previously published boundary-domain-enriched (BD-enriched) genes. These represent genes showing ≥2-fold higher expression in *A. thaliana* organ boundary cells compared to leaf primordia, with statistical significance (adjusted P ≤ 0.001), as determined by cell-type–specific translatome profiling (Tian et al., 2014). BD-enriched genes were significantly overrepresented among HYDRA-up and DCMU-downregulated genes (List A, 964 genes), but not among HYDRA-specific upregulated genes excluding DCMU-downregulated genes (List B, 1901 genes) (Fig. 4C, Supplementary Fig. 5B). This indicates that carbon-dependent transcriptional regulation is strongly associated with the boundary domain.

To determine the spatial location of cell cycle progression, we used the G2/M reporter *CYCB1;2-YFP* line (Iwata et al., 2011). CYCB1;2-YFP fluorescence was detected in the boundary domain upon dissection in HYDRA explants, whereas this region remained largely quiescent in uncut control explants (Fig. 4E). Consistent with transcriptomic data, CYCB1;2-YFP signal was reduced under dark or DCMU conditions and restored by sucrose supplementation (Fig. 4F; Supplementary Fig. 5C), in line with the rescue of shoot regeneration (Fig. 3C, D). Together, these findings demonstrate that carbon availability is required for the transcriptional activation of cell cycle genes in the boundary domain during shoot regeneration in HYDRA.

### Carbon availability promotes cell cycle entry through the RBR1 pathway

Our transcriptomic analyses suggest that carbon deprivation restricts cell cycle progression during shoot regeneration in HYDRA. Carbon resources are linked to cell cycle control, in part through the RBR1 pathway, in which RBR1 inactivation is a key step for activation of cell proliferation programs (Garza-Aguilar et al., 2017; Hirano et al., 2008; Magyar et al., 2012; Riou-Khamlichi et al., 2000) (Fig. 5A). Consistent with this model, multiple *CYCD* genes, including *CYCD3;1*, were upregulated at 30h in a photosynthesis-dependent manner (Fig. 5B). To further assess the involvement of the RBR1 pathway, we compared HYDRA-upregulated genes (List A and List B; Fig. 4C) with published RBR1-, E2FA-, E2FB-, and E2FC-bound targets (Gombos et al., 2023). We observed significant enrichment of RBR1- and E2F-bound genes in our dataset (Supplementary Fig. 6A, B). Importantly, this enrichment was consistently higher in the DCMU-downregulated subset (List A; Fig. 5C), indicating a stronger association of carbon-dependent transcriptional responses with the RBR1 pathway targets.

**Figure 5.**
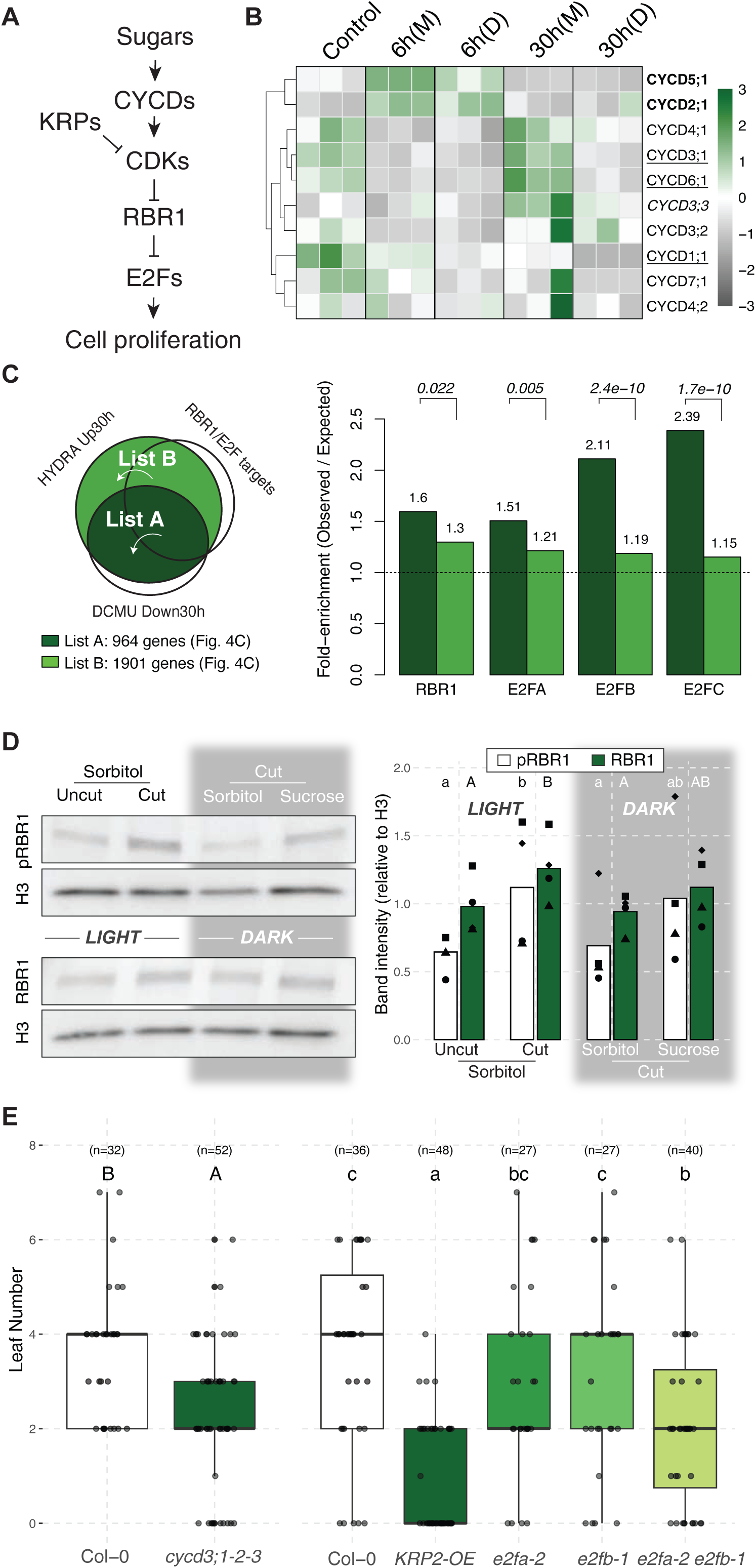
Carbon availability promotes cell cycle entry through the RBR1 pathway in HYDRA. (A) Conceptual model summarizing carbon-dependent cell proliferation.(B) Heatmap of *CYCD* family genes. Significance (FDR < 0.05) is indicated by: **bold (6h),** *italics (30h)*, <u>underlined (DCMU)</u>. (C) Enrichment analyses comparing our dataset with RBR1/E2F-targets (Gombos et al., 2023). P values are indicated. (D) Representative immunoblot (left) of phosphorylated RBR1 (pRBR1), total RBR1 (RBR1), and histone H3 loading control, and their quantification normalized to H3 in HYDRA samples (right). Each dot corresponds to a biological replicate. Bars show average of these values. (E) Quantification of shoot regeneration efficiency in *cycd3;1 cycd3;2 cycd3;3 (cycd3;1-2-3)*, *KRP2-OE*, *e2fa-2*, *e2fb-1*, and *e2fa-2 e2fb-1*. Treatments were applied during the regeneration phase. Boxplots represent the distribution of the data, the center line indicates the median, the box corresponds to the interquartile range (25th to 75th percentile), and whiskers extend to 1.5 × the interquartile range. Different letters indicate statistical significance (P < 0.05). Sample size (n) is indicated.

To identify the molecular processes potentially mediated by the RBR1 pathway in HYDRA, we performed GO term analysis on genes overlapping between HYDRA-upregulated genes and the combined RBR1- and E2F-bound targets (RBR1/E2F targets, 5,820 genes). The 300 genes in List A that overlap with RBR1/E2F targets were strongly enriched for cell cycle and related processes, whereas this enrichment was much less prominent in the other subsets (Supplementary Fig. 6C,D). Notably, these 300 genes were also significantly overrepresented among boundary-domain-enriched genes, unlike the other subsets (Supplementary Fig. 6E), thereby linking carbon-dependent RBR1 pathway activity to the boundary domain. Together, these results suggest that carbon availability promotes cell cycle progression during shoot regeneration in HYDRA, at least in part through RBR1 inactivation in the boundary domain. To test this hypothesis at the biochemical level, we examined phosphorylated RBR1 (pRBR1) levels using a human phospho-RB Ser807/811 antibody validated for detecting *A. thaliana* RBR1 Ser911, a proliferation-associated pRBR1 form depleted from E2F-bound RBR1 complexes (Magyar et al., 2012) (Pettko-Szandtner et al., 2026). pRBR1 levels increased significantly at 30h post-dissection compared to uncut controls (Fig. 5D). This increase was reduced under dark conditions and restored by sucrose supplementation, mirroring the patterns observed for shoot regeneration in HYDRA and CYCB1;2-YFP signal (Figs. 3C, 4F). Total RBR1 levels showed comparatively smaller changes across these conditions. These results indicate that RBR1 phosphorylation depends on carbon availability.

To assess the functional relevance of the RBR1 pathway in HYDRA, we analyzed the *KRP2* overexpression line (*KRP2-OE*), in which CDK activity is inhibited and RBR1 remains in a hypophosphorylated, active state (Magyar et al., 2012; Verkest et al., 2005). As expected, the *KRP2-OE* line showed reduced shoot regeneration in HYDRA (Fig. 5E). Similarly, the *cycd3;1 cycd3;2 cycd3;3* triple mutant (Dewitte et al., 2007), which is expected to reduce RBR1 phosphorylation (Magyar et al., 2012), also exhibited reduced regeneration (Fig. 5E). These results indicate that RBR1 phosphorylation status correlates with shoot regeneration efficiency in HYDRA. To test downstream effectors, we examined *e2fa-2*, *e2fb-1*, and the *e2fa-2 e2fb-1* double mutant (Heyman et al., 2011). While the single *e2fa-2* and *e2fb-1* mutants resembled the wild type, the double *e2fa-2 e2fb-1* mutant showed reduced regeneration (Fig. 5E). To test whether exogenous sugar can bypass the requirement for cell-cycle regulators, we tested *KRP2-OE, cycd3;1 cycd3;2 cycd3;3* and *e2fa-2 e2fb-1* lines under sucrose-supplemented conditions (Supplementary Fig. 6F). These lines remained impaired compared with wild type, indicating that sucrose does not restore regeneration when CYCD3-, E2FA/E2FB-, or CDK-dependent cell-cycle activation is compromised (Supplementary Fig. 6F). Together with the transcriptomic and biochemical evidence, these genetic results support a model in which carbon availability gates cell cycle entry at least partially through the RBR1 pathway during shoot regeneration in HYDRA.

### Meristem regulators function redundantly during shoot regeneration in HYDRA

Carbon-dependent cell cycle activation in the boundary domain accounts for the proliferative response underlying shoot regeneration in HYDRA (Figs. 4, 5). Additionally, the boundary domain serves as the source for shoot regeneration capacity (Fig. 2). However, what makes this domain special such that carbon-dependent cell cycle activation specifically in this domain leads to shoot meristem formation remains unclear. Boundary domains are characterized by strong expression of meristem regulators including *STM*, *LAS*, and *CUCs* (Aida et al., 1997; Aida et al., 1999; Greb et al., 2003; Hibara et al., 2006; Keller et al., 2006; Long and Barton, 2000; Raatz et al., 2011; Rast and Simon, 2008; Tian et al., 2014). We therefore examined the expression of the four genes that are redundantly required for cotyledon boundary formation, *STM*, *LAS*, *BLR*, and *KNAT6*, as well as their upstream regulators *CUCs* (Aida et al., 2020; Belles-Boix et al., 2006). Notably, none of these genes were transcriptionally induced during HYDRA (Fig. 6A; Supplementary Fig. 7A). Instead, these transcripts were already present in uncut controls, unlike *WUS* and *CLV3,* which were undetectable before dissection and became induced only during regeneration (Fig. 4B). Consistent with this observation, *STM* reporter signal was strongly detected in the boundary domain prior to dissection (Fig. 6B), overlapping spatially with the subsequent *CYCB1;2*, *WUS* and *CLV3* expression domains (Figs. 1D, 4F). These observations suggest that pre-existing expression of boundary domain regulators underlies the specific capacity of this domain to support shoot regeneration in HYDRA.

**Figure 6.**
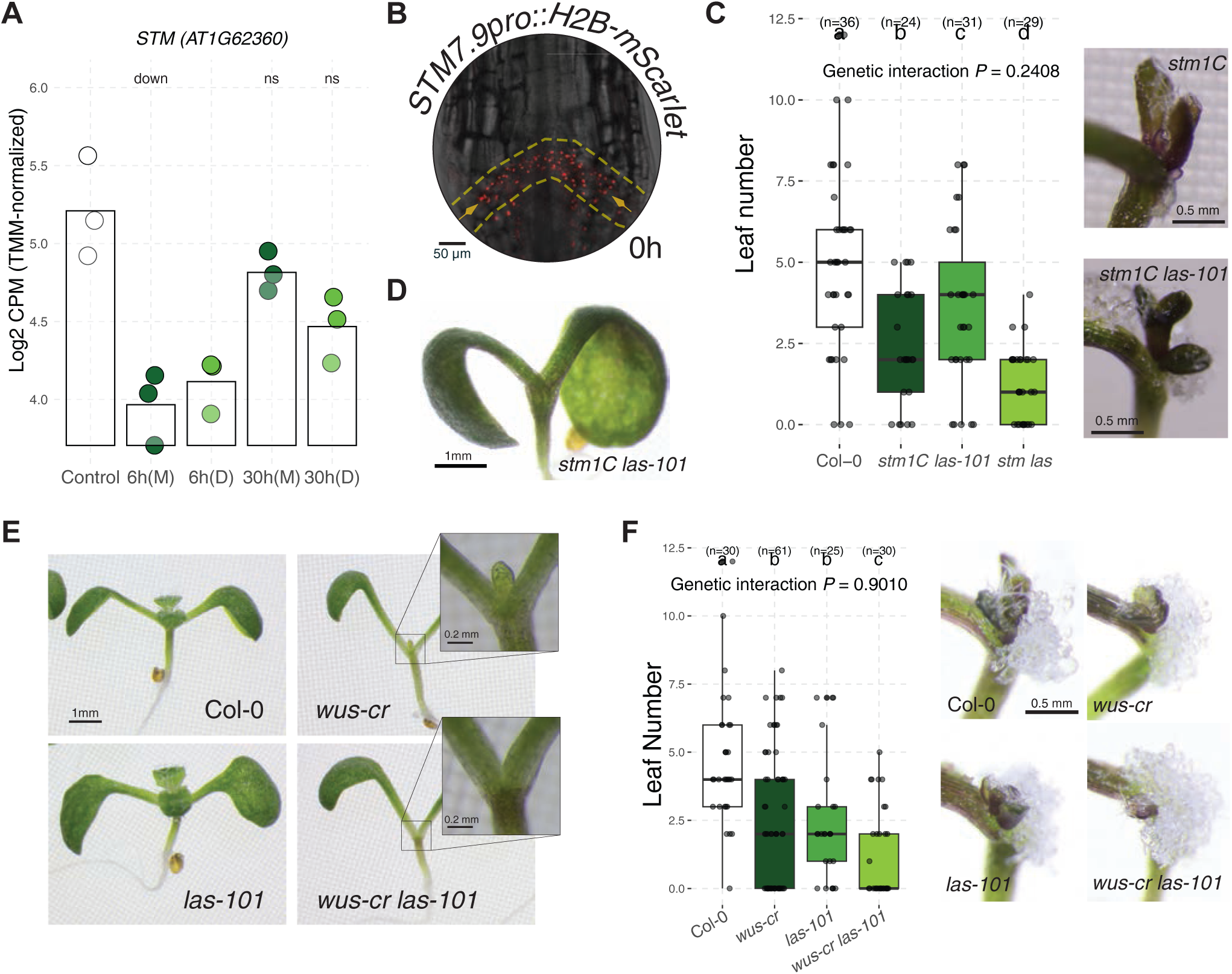
Meristem regulators function redundantly during shoot regeneration in HYDRA. (A) Expression of *STM* in our dataset (ns: not significant, down: downregulated FDR <0.05). (B) STM7.9pro::H2B-mScarlet signal in HYDRA at 0h. The region between yellow lines marks the boundary domain. Image is a single optical section. (C) Quantification (left) and representative images (right) of shoot regeneration in *stm1C, las-101,* and *stm1C las-101*. (D) SAM phenotype in *stm1C las-101* mutant. (E) SAM phenotypes in *wus-cr, las-101,* and *wus-cr las-101*. (F) Quantification (left) and representative images (right) of shoot regeneration in *wus-cr, las-101,* and *wus-cr las-101*. Media were supplemented with sucrose. Boxplots represent the distribution of the data, the center line indicates the median, the box corresponds to the interquartile range (25th to 75th percentile), and whiskers extend to 1.5 × the interquartile range. Different letters indicate statistical significance (P < 0.05). Sample size (n) is indicated.

To test their functional importance, we analyzed boundary domain mutants in HYDRA. The *stm1C* mutant (Takano et al., 2010) showed a strong reduction in shoot regeneration, although residual regeneration persisted, suggesting redundancy (Fig. 6C). Consistent with this, the *las-101* mutant (Hibara et al., 2006) displayed reduced regeneration, while the *stm1C las-101* double mutant (Aida et al., 2020) exhibited a stronger defect (Fig. 6C). Nevertheless, regeneration was not completely abolished, indicating further redundancy among other factors. This phenotype is also reminiscent of cotyledon boundary domain formation, where loss of these four genes leads to a progressive impairment of boundary domain formation (Aida et al., 2020). Because *CUCs* act upstream of these four genes and high-order *cuc* mutants completely lack cotyledon boundary domain formation (Aida et al., 2020; Hasson et al., 2011; Hibara et al., 2006), we tested these mutants in HYDRA to overcome the redundancy among these four downstream factors. While single *cuc2* alleles regenerated like WT, higher order *cuc* mutants involving *cuc2* alleles abolished regeneration (Supplementary Fig. 7B). However, the *CUC2* reporter (Temman et al., 2023) was not detectable in the boundary domain immediately after dissection unlike *STM* (Supplementary Fig. 7C). This supports that the requirement for CUCs likely reflects their established role upstream of these four factors.

In addition to their roles in boundary domain formation, these factors are also key regulators of primary SAM establishment (Aida et al., 2020; Endrizzi et al., 1996; Hibara et al., 2006). Consistent with previous reports, the *stm1C las-101* mutant completely lacked a primary SAM structure under our conditions (Fig. 6D). Loss of *WUS* is also known to cause severely impaired SAM function (Laux et al., 1996). Accordingly, the *wus-cr* allele showed strongly impaired SAM development, although a small residual SAM structure remained visible (Fig. 6E; Supplementary Fig. 7D). In line with this, *wus-cr* showed a significant but partial reduction in shoot regeneration (Fig. 6F). However, shoot regeneration still occurred in *wus-cr*, so we asked whether boundary domain regulators could account for this residual regenerative capacity. Given their established roles in SAM development, these factors were strong candidates to function redundantly with *WUS* during shoot regeneration. The *wus-cr las-101* double mutant exhibited a stronger defect in SAM establishment than *wus-cr* alone (Fig. 6E), supporting the idea of redundancy between WUS and LAS during SAM formation. In line with this, *wus-cr las-101* showed a further reduction in shoot regeneration compared to either single mutant (Fig. 6F), suggesting that *WUS* contributes to de novo meristem establishment in HYDRA together with *LAS*. Taken together, these results indicate that boundary domain-associated meristem regulators function redundantly to enable shoot regeneration in HYDRA, linking pre-existing boundary domain identity to de novo meristem formation upon carbon-dependent proliferative activation.

### Carbon availability overrides apical dominance enabling shoot regeneration

Our results establish that shoot regeneration in HYDRA is spatially restricted to the boundary domain and depends on carbon-driven cell cycle activation following removal of apical dominance. However, the mechanism by which the removal of apical dominance is perceived specifically in the boundary domain remains unclear. In axillary bud activation, apical dominance is largely mediated by preferential carbon allocation to the SAM, which acts as a strong sink (Schneider et al., 2019). Removal of the SAM redistributes carbon resources, increasing their availability to the axillary buds and enabling their outgrowth (Fichtner et al., 2021; Fichtner et al., 2017; Mason et al., 2014). We therefore hypothesized that, in HYDRA, carbon availability functions as the key cue linking the removal of apical dominance to initiation of shoot regeneration. Supporting this idea, genes associated with carbohydrate transport (GO:0008643; 64/146 genes) were enriched among the HYDRA-upregulated genes (Fig. 7A). Significantly upregulated genes included members of the *SUCROSE TRANSPORTER* (*SUC*), *SUGARS WILL EVENTUALLY BE EXPORTED TRANSPORTER* (*SWEET*), *SUGAR TRANSPORT PROTEIN* (*STP*), and *EARLY RESPONSE TO DEHYDRATION 6-LIKE* (*ERDL*) families, which mediate long-distance sucrose transport, cellular import and export, and mobilization from intracellular stores, respectively (Lei et al., 2025). Sucrose is the main form of long-distance sugar transport and can be hydrolyzed into glucose and fructose by CELL WALL INVERTASEs (CWIs), which are considered markers of active sink tissues (Proels and Huckelhoven, 2014). Consistent with this, we observed an approximately 40-fold increase in *CWI1* expression 6h after dissection (Fig. 7B). We also quantified sucrose, glucose, and fructose levels in boundary-enriched samples (as in Fig. 4A). To obtain sufficient material for sugar quantification, we used *Brassica rapa* seedlings, which regenerate shoots similarly to *Arabidopsis thaliana* in HYDRA (Supplementary Fig. 1B), supporting their use for these measurements. Sucrose levels remained unchanged in cut samples compared with uncut controls (Fig. 7C; Supplementary Fig. 8A). In contrast, the combined levels of glucose and fructose increased significantly at 18h and 24h after dissection (Fig. 7C; Supplementary Fig. 8A). This is consistent with the upregulation of sucrose transport genes and *CWI1*, and indicates that carbon availability increases in the boundary domain after dissection.

**Figure 7.**
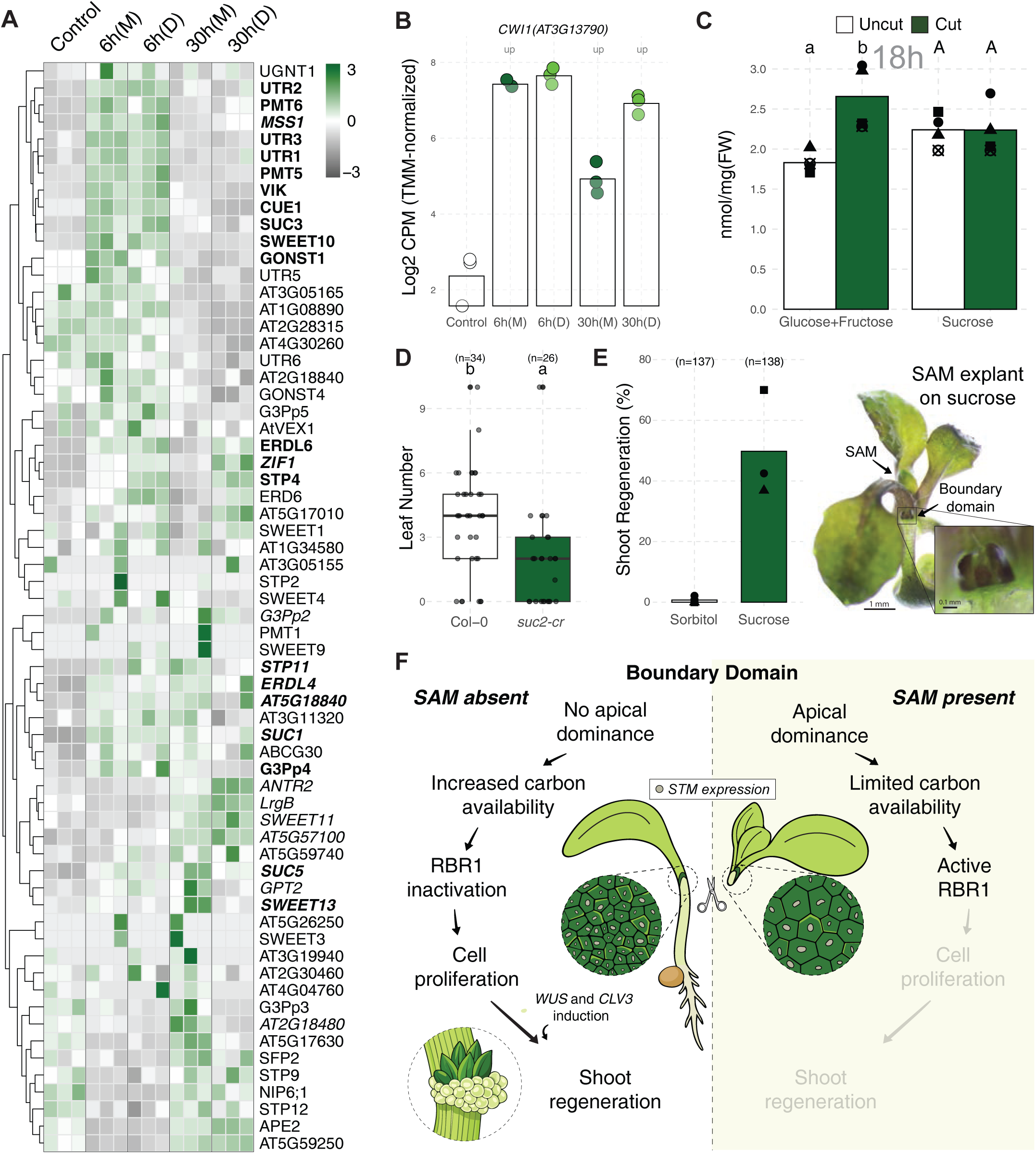
Carbon availability overrides apical dominance to enable shoot regeneration. (A) Heatmap of genes from Carbohydrate Transport (GO: 0008643) found among HYDRA-upregulated genes. **Bold (6h)** and *italics (30h)* indicate significance (FDR < 0.05). (B) Expression of *CWI1*. up: upregulated (FDR<0.05). (C) Sugar quantification in *B. rapa* boundary-enriched samples 18h after dissection in comparison to control (uncut). (D) Shoot regeneration in a sucrose transport mutant. Boxplots represent the distribution of the data, the center line indicates the median, the box corresponds to the interquartile range (25th to 75th percentile), and whiskers extend to 1.5 × the interquartile range. Different letters indicate statistical significance (P < 0.05). Sample size (n) is indicated. (E) Shoot regeneration in SAM explants. Total sample size (n) is indicated. (C, E) Each dot corresponds to a biological replicate. Bars show average of these values. (F) Working model for shoot regeneration in HYDRA. Upon SAM removal, increased carbon availability promotes cell proliferation in otherwise quiescent boundary domain via RBR1 inactivation. This is accompanied by induction of meristem genes (e.g., *WUS* and *CLV3*), ultimately enabling shoot regeneration in HYDRA.

To assess functional relevance of sucrose transport, we generated a weak allele *suc2-cr,* targeting the last exon of *SUC2* that encodes a key long-distance sucrose transporter (Supplementary Fig. 8B). We validated *suc2-cr* by increased starch accumulation in cotyledons, consistent with impaired sucrose transport (Supplementary Fig. 8C). The *suc2-cr* mutant showed significantly reduced regeneration (Fig. 7D), indicating that sucrose transport is functionally important during shoot regeneration in HYDRA. If carbon availability serves as the initiation cue, exogenous sugars should bypass apical dominance in SAM explants that completely fail to regenerate shoots (Fig. 2D). Therefore, we treated SAM explants with exogenous sucrose. Sucrose induced shoot regeneration in a substantial fraction of SAM explants, whereas sorbitol had no effect (Fig. 7E), demonstrating that increased carbon availability can bypass SAM-dependent suppression of regeneration at the boundary domain. Together, these findings indicate that carbon availability also acts as a key downstream effector of apical dominance removal, enabling initiation of shoot regeneration in HYDRA.

## DISCUSSION

In this study, we establish HYDRA as a new shoot regeneration system for studying hormone-autonomous regeneration in *Arabidopsis thaliana*. Our findings support a two-component model for shoot regeneration in HYDRA (Fig. 7F). First, photosynthesis-derived carbon drives cell cycle activation in the boundary domain, with the RBR1 pathway acting as a mediator of this response. Second, meristem marker genes, *WUS* and *CLV3*, are induced in the boundary domain independently of carbon availability. Both processes are required because carbon deprivation blocks regeneration despite meristem marker induction, whereas perturbation of the boundary domain, its regulators, or *WUS* reduces regeneration despite sufficient carbon. Pre-existing expression of boundary-associated regulators, as shown for *STM*, defines the boundary domain as the site where these two requirements converge. Together, carbon-dependent proliferation and carbon-independent meristem gene induction represent central requirements for hormone-autonomous shoot regeneration in HYDRA.

Although shoot regeneration in HYDRA differs from axillary meristem (AM) initiation in that *A. thaliana* cotyledons do not form AMs (Grbic and Bleecker, 2000) and HYDRA frequently produces multiple shoots unlike AM initiation, the two systems nevertheless share common features at three levels. First, boundary domains are associated with the spatial origin in both systems. Second, these domains harbor slow-dividing cells with pre-existing expression of key meristem regulators, including *STM* (Aida and Tasaka, 2006; Balkunde et al., 2017; Cao et al., 2020; Guo et al., 2025; Hepworth and Pautot, 2015; Keller et al., 2006; Long and Barton, 2000; Shi et al., 2016; Wang and Jiao, 2018; Zhang et al., 2018). Third, meristem formation in both systems depends on proliferative activation, indicating that pre-existing meristem regulator expression must be coupled to cell cycle entry to generate a functional meristem. Consistently, *STM* expression alone is not sufficient to trigger cell division (Gallois et al., 2002; Guo et al., 2025; Shi et al., 2016). In HYDRA, reduced regeneration in the *cycd3;1-2-3* mutant and the *KRP2-OE* line supports that this activation is driven by RBR1 inactivation. A similar mechanism likely operates during AM initiation, where *CYCD3;1* promotes and *KRP4* inhibits AM formation when expressed in the boundary domain (Guo et al., 2025). Together, these observations support a conserved mechanism in which boundary domains provide shoot competence, and proliferative activation through RBR1 inactivation enables shoot meristem formation.

Shoot regeneration in HYDRA also parallels the activation of axillary bud outgrowth in terms of the initiation signal. Apical dominance is largely mediated by carbon allocation, with SAM functioning as a major sink that limits resource availability to axillary buds (Fichtner et al., 2021; Fichtner et al., 2017; Mason et al., 2014). SAM removal redistributes carbon and enables bud activation. Similarly, in HYDRA, SAM removal is required for regeneration and is accompanied by transcriptional activation of sugar transport, increased hexose content in the boundary domain, and a functional requirement for sugar transport. Furthermore, exogenous sugar application is sufficient to overcome apical dominance and induce shoot formation, indicating that carbon availability acts also as an initiation cue. These observations support that carbon availability functions at two levels; (i) it contributes to the establishment of apical dominance and acts as an initiation cue upon SAM removal, and (ii) promotes cell cycle progression leading to meristem formation.

In line with this framework, although auxin contributes to shoot regeneration in HYDRA, the system tolerates substantial perturbations in auxin pathway components. This may be because carbon availability and pre-existing boundary domain identity in HYDRA largely fulfill the proliferative and competence-related roles of auxin, respectively. However, given the crosstalk between sugars and auxin in various contexts (Kiba et al., 2019; Kircher and Schopfer, 2023; Mishra et al., 2009; Paulišić et al., 2026; Sairanen et al., 2012), the potential contribution of carbon availability to this robustness, as well as the extent of this redundancy, remain to be further investigated.

Although sugars can influence SAM regulators in established meristems over longer timescales (Lopes et al., 2024; Pfeiffer et al., 2016), carbon availability does not regulate meristem gene induction at early stages of HYDRA. Together, these observations support a distinction between early meristem gene induction and the formation of a visible, functional meristem. While *WUS* and *CLV3* transcripts are induced under carbon-limited conditions, organized meristem formation requires sufficient cell proliferation, which depends on carbon availability. Thus, carbon does not appear to directly initiate early meristem gene expression but is required to convert this transcriptional state into a functional shoot meristem through proliferative growth, potentially explaining how sugars influence meristem regulators over longer developmental timescales (Lopes et al., 2024; Pfeiffer et al., 2016). The rapid carbon-independent activation of *WUS* may instead reflect a permissive transcriptional or epigenetic environment within the boundary domain. Consistent with this idea, boundary-associated regulators such as *STM* are already expressed before regeneration and shown to directly regulate SAM-associated genes, including *CLV3* (Balkunde et al., 2017; Brand et al., 2002; Hepworth and Pautot, 2015; Lechon et al., 2025; Scofield et al., 2018; Spinelli et al., 2011). In addition, cytokinin remains functionally required in HYDRA, despite the high redundancy in cytokinin signaling. Cytokinin is a well-established regulator of *WUS* expression, and this regulation depends on a permissive epigenetic environment during AM initiation (Iwase et al., 2017; Meng et al., 2017; Smeringai et al., 2023; Wang et al., 2017; Zhang et al., 2017). Notably, epigenetic regulation is closely linked to cell proliferation in several developmental processes including during AM initiation (Guo et al., 2025; Liu et al., 2018; Rahni et al., 2026; Williams et al., 2003). Finally, wound-induced transcription factors provide additional inputs into *WUS* activation and epigenetic regulation (Ikeda et al., 2006; Ikeda et al., 2021; Iwase et al., 2017; Iwase et al., 2026). Dissecting the relative contributions of these mechanisms and defining the sequence of events linking boundary domain to *WUS* activation will require further investigation.

In conclusion, HYDRA reveals that *Arabidopsis thaliana* possesses an inherent capacity for hormone-autonomous shoot regeneration. This autonomy, together with the robustness, accessibility, and cross-species applicability of the assay, makes it a valuable tool for investigating regulatory mechanisms of shoot regeneration. These features position HYDRA as a complementary system to classical regeneration assays, enabling further dissection of the integration of metabolic, developmental, and environmental cues in shoot regeneration.

## MATERIALS AND METHODS

### Materials and growth conditions

Complete lists of plant materials, chemical materials, oligonucleotides, and sgRNAs are provided in Supplementary Table 3 to 6 respectively. Seeds were surface-sterilized in 70% (v/v) ethanol containing 0.05% (v/v) Triton X-100, followed by 100% (v/v) ethanol for 10 min each. Seeds were stratified at 4 °C for 3 days in dark and germinated vertically on Murashige and Skoog (MS) medium supplemented with 6 g/L (w/v) gelrite and 10 g/L sucrose for homogeneous growth on a nylon mesh. SIM is B5 media supplemented with B5 vitamins, MES (0.5 g/L), Glucose (20g/L), 2iP (0.5 mg/L), IBA (0.1 mg/L), and gelrite 6 g/L (w/v). Seven-day-old seedlings were dissected under a binocular microscope in a clean bench and transferred to MS media with or without sugar supplementation and/or specific light regimes, as indicated in the figure legends. Unless otherwise specified, seedlings were grown under continuous white light (WL, ∼40 μmol/m² s¹) provided by fluorescent tubes (Mitsubishi Osram FL40SSW37), at 22 °C. Blue (400-500 nm) and red (R, 600-700 nm) light were applied using LED sources (∼10 μmol/m² s¹). WL supplemented with far-red (FR, 700-780 nm) LEDs (95 μmol/m² s¹), resulting in near R/FR ratios of 1.96 (WL) and 0.04 (WL+FR) (Ince and Galvao, 2021) was used as shade treatment. For sugar rescue, sorbitol (15.96 g/L; same molarity as sucrose), sucrose (30 g/L), and glucose (31.6 g/L; energy-equivalent to sucrose) were added to MS media. Sucrose supplementation to SAM explants was done by liquid medium with a nylon mesh to ensure direct exposure of the boundary domain. Because homozygous *pds3-cr* and *wus-cr* mutations are not viable, experiments were performed using pooled T2 progeny from multiple independent lines selected based on bleaching (*pds3-cr*) and no SAM (*wus-cr*) phenotype. Disruptive mutations were confirmed by sequencing in representative lines.

### Shoot regeneration quantification

Shoot regeneration was quantified at 7 days post-dissection using two different approaches, depending on the experimental question. For binary assessments of regeneration capacity, we scored regeneration frequency across samples. For quantitative comparisons of regeneration strength among treatments or genotypes, we quantified the number of regenerated outermost leaves (Supplementary Fig. 1F).

### Constructs and cloning

For *suc2-cr*, a single sgRNA was cloned into *pHEE401E* (Wang et al., 2015). For *wus-cr* and *pds3-cr*, the intron-containing *zCas9* (Grutzner et al., 2021) was cloned into XbaI-SacI digested *pEC1.2/EC1.1_spCas9_TagRFP* (Christian Fankhauser Lab, Addgene plasmid # 161934) generating *pYI061*. PCR-amplified *sgRNA1::scaffold::tRNA::sgRNA2* (*pGTR* template, (Xie et al., 2015) was cloned into *pYI061*. *STM7.9pro::H2B-mScarlet* was constructed using PCR-amplified CDS of H2B (AT5G22880) and mScarlet (Bindels et al., 2017) separated by GGGGGATCTGGCGGT linker, in between 7.9-kb upstream (36,012–28,120, BAC F24O1), 0.8-kb downstream (25,195–24,407, BAC F24O1) of *STM*, into *pH7m34GW* (Karimi et al., 2005). All constructs were verified by Sanger sequencing and introduced into Col-0 via *Agrobacterium tumefaciens* GV3101 using the floral dipping (Clough and Bent, 1998).

### RNA isolation, library preparation, and RNA sequencing

L*er* seedlings were dissected as in Fig. 4A, immediately frozen in liquid nitrogen. Total RNA integrity and library quality were assessed using a Fragment Analyzer (Agilent Technologies) and a Qubit fluorometer (ThermoFisher Scientific). Sequencing was performed by Macrogen Japan on an Illumina NovaSeq X Plus platform using paired-end 150 bp reads. Raw sequencing data were demultiplexed using bcl2fastq Conversion Software (Illumina).

### RNA-seq data analysis

Raw reads were quality-checked using FastQC and trimmed using fastp (Andrews S. 2010; Chen et al., 2018), with quality filtering (q ≥ 15) and length filtering (≥5 bp), resulting in high-quality reads with Q20 >90% and Q30 >80%. Clean reads were aligned to the *Arabidopsis thaliana* reference genome (TAIR10.56) using STAR (Dobin et al., 2013). Gene-level counts were obtained using featureCounts. For downstream analyses, edgeR and limma (Ritchie et al., 2015; Robinson et al., 2010) in R (http://cran.r-project.org/) were used. Genes with a total count >1 across all samples were retained, and normalization was performed using the TMM method (Robinson and Oshlack, 2010). Expression values were calculated as TMM-normalized CPM, and log2-transformed CPM values were generated with a prior count of 1 (Law et al., 2014). PCA was performed using prcomp, and hierarchical clustering was conducted using Euclidean distance and complete linkage on Z-score–transformed data. Differential expression analysis was performed using the limma-trend workflow with empirical Bayes moderation (Smyth, 2004), and P values were adjusted (Benjamini and Hochberg, 1995). Comparisons included cut versus uncut (at 6h and 30h) and mock versus DCMU (at 30h) (FDR < 0.05). Heatmaps were generated using pheatmap with clustering applied to genes and was based on control and mock samples, while all conditions were displayed. The g:Profiler (g:GOSt, version *e114_eg62_p19_fa3a7d2c*) (Raudvere et al., 2019) was used for Gene Ontology analysis with default parameters and to obtain gene IDs for given GO terms used in boxplot representations.

### Western blot analysis

Total protein extracts were prepared from dissected tissues (Fig. 4A) as described previously (Fiorucci et al., 2019; Ince et al., 2022). Samples were ground in Final Sample Buffer (FSB), incubated at 95 °C for 7 min, and centrifuged for 10 min at 4 °C. Proteins were separated on precast gels and transferred to PVDF membranes using a Turbo transfer system. After blocking in 5% (w/v) milk for 1h at RT, membranes were incubated with primary and secondary antibodies were overnight at 4 °C and 1h at RT, respectively. Signals were detected on an ImageQuant LAS 4000 (GE Healthcare). Western blot analyses were performed with four biological replicates and quantified using ImageJ.

### Microscopy

mScarlet and dsRed7 (excitation 558 nm, emission mScarlet; 568-650 nm, emission dsRed7; 573–594 nm), YFP (excitation 514 nm, emission 524–560 nm), and GFP (excitation 488 nm, emission *CUC2g-GFP*; 498–526 nm, *gWUS-3xGFP* 500-545 nm) signals were detected with ×20 objective (NA 0.75, DIC) (Leica confocal microscope, TCS SP8 WLL FALCON). For *CYCB1;2*-YFP, image stacks were acquired until the signal was no longer detectable (pinhole 1 au; 40-50 μm section; 5 μm interval). YFP-expressing cells were manually counted along the boundary domain, excluding surrounding cells from maximum projections.

### Sugar quantification

Glucose, fructose, and sucrose concentrations were measured using an enzyme-based assay as described previously (Cross et al., 2006), with minor modifications. Boundary-enriched tissues from 7d old Brassica rapa seedlings were collected, weighed, flash-frozen in liquid nitrogen, and ground using a TissueLyzer (Qiagen). Ethanolic extracts were prepared by two sequential extractions with 80% ethanol/10 mM HEPES-KOH, pH 7.0, at 80°C for 20 min, followed by a third extraction with 50% ethanol/10 mM HEPES-KOH, pH 7.0. Pooled extracts (650µl) were dried in a SpeedVac concentrator and reconstituted in 200µl water before analysis. 40µl of each extract were assayed in triplicates in 96-well plates in assay buffer containing HEPES-KOH, 4mg/ml ATP, 2.4mg/ml NADP disodium salt, and 3.5U/ml glucose-6-phosphate dehydrogenase. Absorbance at 340 nm was measured after sequential addition of hexokinase, phosphoglucose isomerase, and invertase to determine glucose, fructose, and sucrose, respectively. Sugar concentrations were calculated using standards measured on the same plate and normalized to fresh weight.

### Iodine staining

Iodine staining was performed as described previously (de Wit et al., 2018). Seedlings were heated in 80% ethanol until cleared of chlorophyll, and subsequently stained in Lugol solution for 5 min and rinsed in distilled water for 2 min.

### Statistical analysis

Statistical analyses were performed in R. Data were reshaped into long format, and experimental groups were defined by combining relevant variables. For western blot and sugar content analyses, linear mixed-effects models were fitted using the lmer function (lme4 package) with condition as a fixed effect and biological replicate as a random intercept (Value ∼ Condition + (1 | Biorep)). Significance was assessed using ANOVA (lmerTest package). Shoot regeneration phenotypes (leaf number per seedling) were analyzed using generalized linear models (GLM) with a negative binomial (NB) distribution (glm.nb, MASS package) to account for overdispersion. For interaction analyses, a 2×2 factorial design was implemented and models included both main effects and interaction terms (Value ∼ Factor1 * Factor2), with significance assessed using Type II ANOVA (car package). Gene set analyses used Z-score–transformed expression values, with statistical testing by one-way ANOVA followed by Tukey’s HSD (P < 0.05). For enrichment analyses, overlap between gene sets was assessed using Fisher’s exact test based on contingency tables, and fold-enrichment was calculated relative to background gene frequencies. Estimated marginal means and pairwise comparisons were calculated using the emmeans package (type = “response” where applicable). Compact letter displays were generated to summarize statistical groupings at α = 0.05. Data were visualized using ggplot2.

### Data availability

The RNA-seq data generated in this study are accessible through accession number GSE333850 in NCBI’s Gene Expression Omnibus (GEO).

## Author contributions

Y.C.I.: conceptualization, formal analysis, data curation, methodology, investigation, validation, visualization, resources, funding acquisition, supervision, writing—original draft, and writing—review and editing. A.T.: investigation and validation. A.I.: investigation, resources, and supervision. J.K.: investigation, resources, and methodology. A.C.: resources. M.A.: resources. L.D.V.: resources, and writing—review and editing. K.S.: conceptualization, project administration, resources, funding acquisition, supervision, and writing—review and editing. All authors read and approved the manuscript.

## Acknowledgements

We thank Mariko Mouri and Noriko Doi for technical assistance. We are grateful to Yu Chen, Dongbo Shi, Max Fishman, and Ken Shirasu (RIKEN), Christian Fankhauser and Edward Farmer (University of Lausanne), Sachihiro Matsunaga (University of Tokyo), Momoko Ikeuchi (Nara Institute of Science and Technology), Lin Xu (CAS CEMPS), Markéta Pernisová (Masaryk University), and Lena Hyvärinen and Felix Kessler (University of Neuchâtel) for providing materials and resources. We thank Dongbo Shi (RIKEN) for helpful discussions and comments on the manuscript.

## SUPPLEMENTARY INFORMATION

**Supplementary Figure 1.**
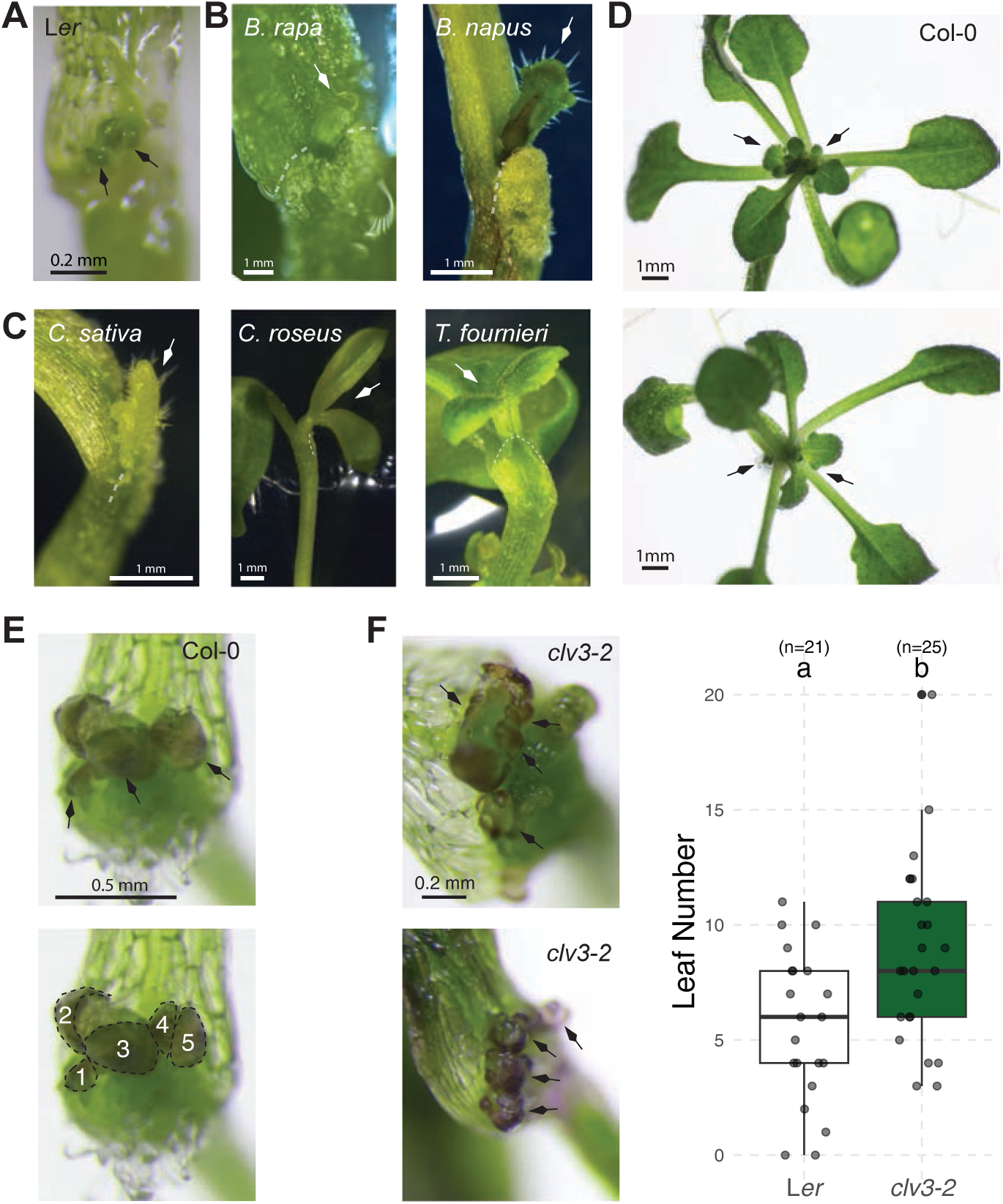
HYDRA is a de novo regeneration assay applicable to multiple species (Related to Figure 1). (A) Shoot regeneration in HYDRA explants cultured on filter paper moistened with water only. (B) HYDRA shoot regeneration in other *Brassicaceae* species: *Brassica rapa, Brassica napus*. (C) HYDRA shoot regeneration in other example dicot species: *Camelina sativa, Catharanthus roseus, Torenia fournieri*. (D) Example of multiple regenerated shoots with normal leaf morphology continuing growth (top) and one shoot becomes dominant while others remain dormant (bottom). (E) Scoring for shoot regeneration quantification. Leaves were counted at early emergence to avoid secondary leaf formation from the same meristem. Leaves outside the field of view are not shown; seedlings were examined from multiple angles to ensure accurate counting of all regenerated leaves. (F) Examples of multiple leaves regenerating from an enlarged united meristem (top) or multiple unconnected meristems (bottom) and shoot quantification (right) in *clv3-2* mutant. Individual data points are shown as dots. Boxplots represent the distribution of the data, the center line indicates the median, the box corresponds to the interquartile range (25th to 75th percentile), and whiskers extend to 1.5 × the interquartile range. Different letters indicate statistical significance (P<0.05). Sample size (n) is indicated. Arrow heads mark regenerating structures. Dashed lines indicate the boundary between cotyledon and hypocotyl.

**Supplementary Figure 2.**
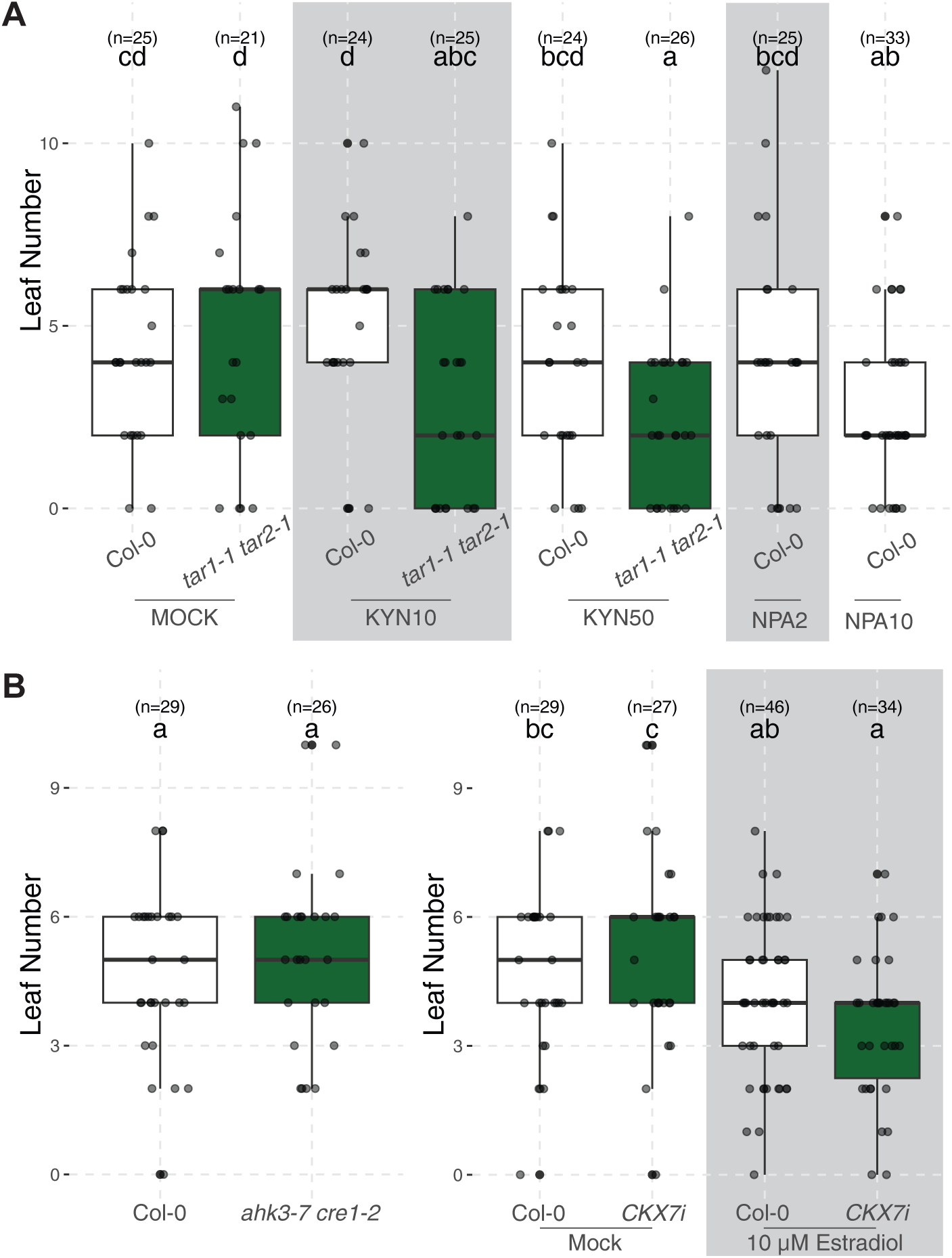
Shoot regeneration in HYDRA tolerates substantial perturbations in endogenous auxin and cytokinin pathways. (Related to Figure 1). (A) Quantification of shoot regeneration in auxin biosynthesis mutants and chemical inhibitors for auxin biosynthesis (KYN: L-Kynurenine) or auxin transport (NPA: N-1-naphthylphthalamic acid). Numbers indicate molarity in µ scale (e.g. KYN10: L-Kynurenine 10 µM). (B) Quantification shoot regeneration in cytokinin receptor mutant (*ahk3-7 cre1-2*) and inducible *CKX7* overexpression (*CKX7i*) line. Media were supplemented with sucrose. Boxplots represent the distribution of the data, the center line indicates the median, the box corresponds to the interquartile range (25th to 75th percentile), and whiskers extend to 1.5 × the interquartile range. Different letters indicate statistical significance (P < 0.05). Sample size (n) is indicated.

**Supplementary Figure 3.**
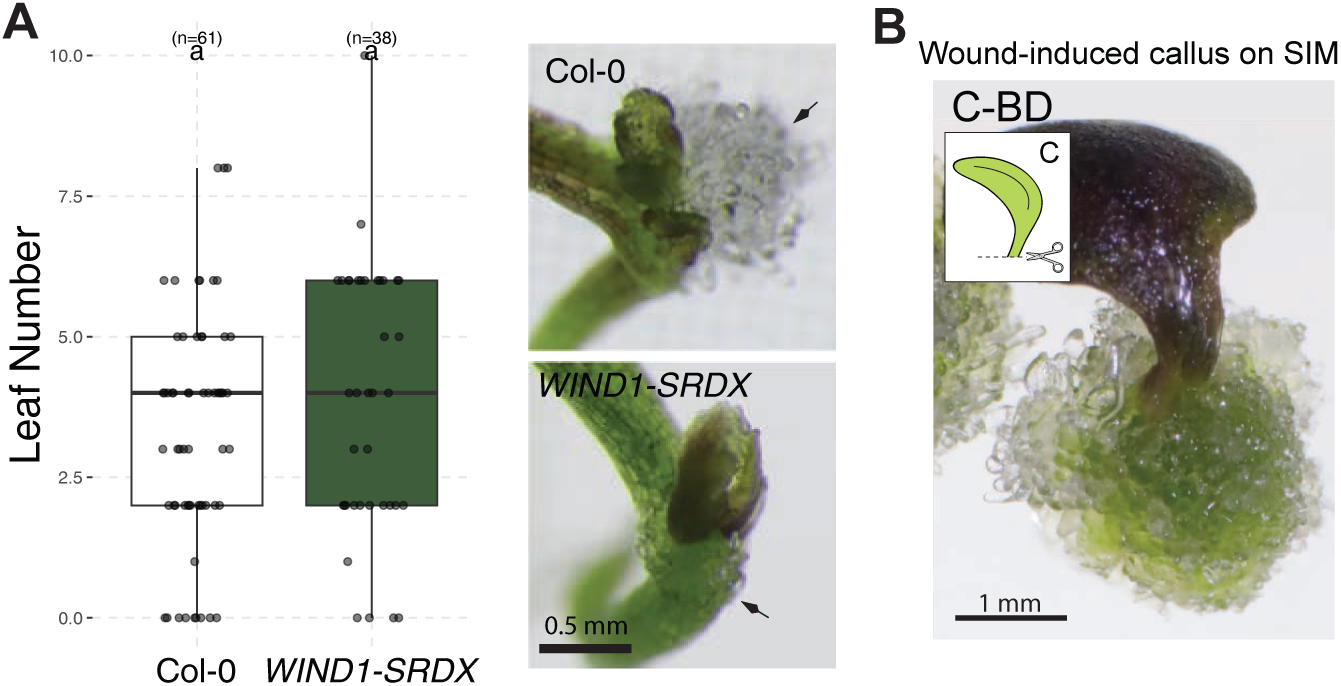
Wound-induced callus (WIC) formation is not required for shoot regeneration in HYDRA (Related to Figure 2). (A) Quantification (left) and representative images (right) of shoot regeneration in *WIND1-SRDX* line. Arrowheads point presence or absence of WIC. Boxplots represent the distribution of the data, the center line indicates the median, the box corresponds to the interquartile range (25th to 75th percentile), and whiskers extend to 1.5 × the interquartile range. Different letters indicate statistical significance (P < 0.05). Sample size (n) is indicated. (B) Representative image of WIC on cotyledon explant without boundary domain (see Fig. 2) after transfer to SIM for 14 days.

**Supplementary Figure 4.**
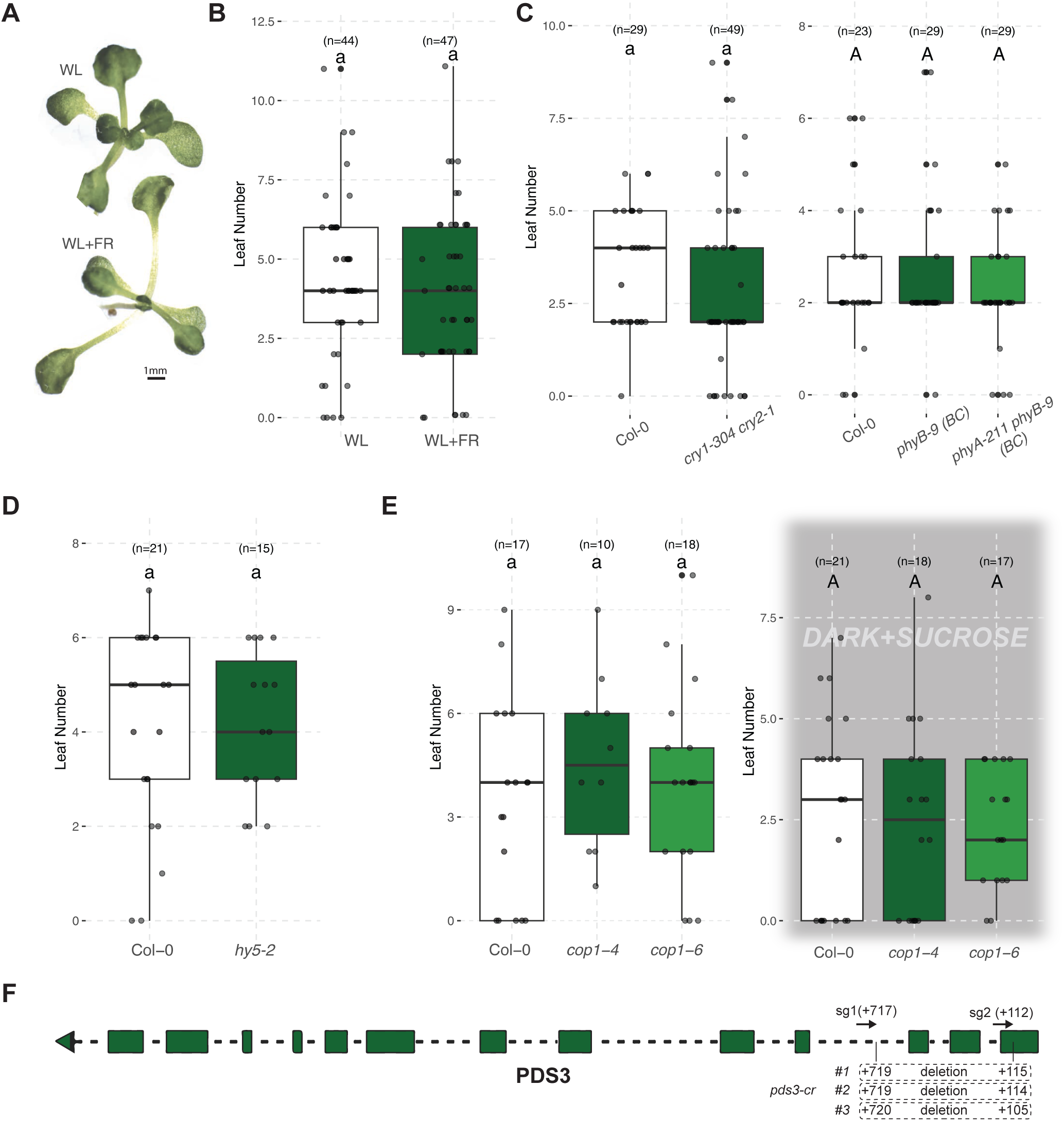
Major light signaling components do not contribute to shoot regeneration in HYDRA (Related to Figure 3). (A) Morphology of uncut seedlings grown under white light (WL) or WL supplemented with far-red (WL+FR), showing WL+FR-induced shade-avoidance response with elongated petioles. (B) Shoot regeneration under WL and WL+FR during the regeneration phase. (C) Shoot regeneration in photoreceptor mutants. To minimize initial growth defects, *cry1-304 cry2-1* was grown under red light, whereas *phyB-9 (BC)* and *phyA-211 phyB-9 (BC)* were grown under blue light together with their corresponding WT control prior to dissection. Regeneration assays were performed under white light. (D) Shoot regeneration in *hy5-2*. (E) Shoot regeneration in *cop1* mutants with light (left) and dark supplied with sucrose (right). (F) Genomic structure of *PDS3* showing sgRNA target sites and sequencing confirmation of induced mutations. Dashed lines indicate introns, boxes exons, and arrowheads PAM sequences. Locations (bp) are given relative to start codon ATG. Boxplots represent the distribution of the data, the center line indicates the median, the box corresponds to the interquartile range (25th to 75th percentile), and whiskers extend to 1.5 × the interquartile range. Different letters indicate statistical significance (P < 0.05). Sample size (n) is indicated.

**Supplementary Figure 5.**
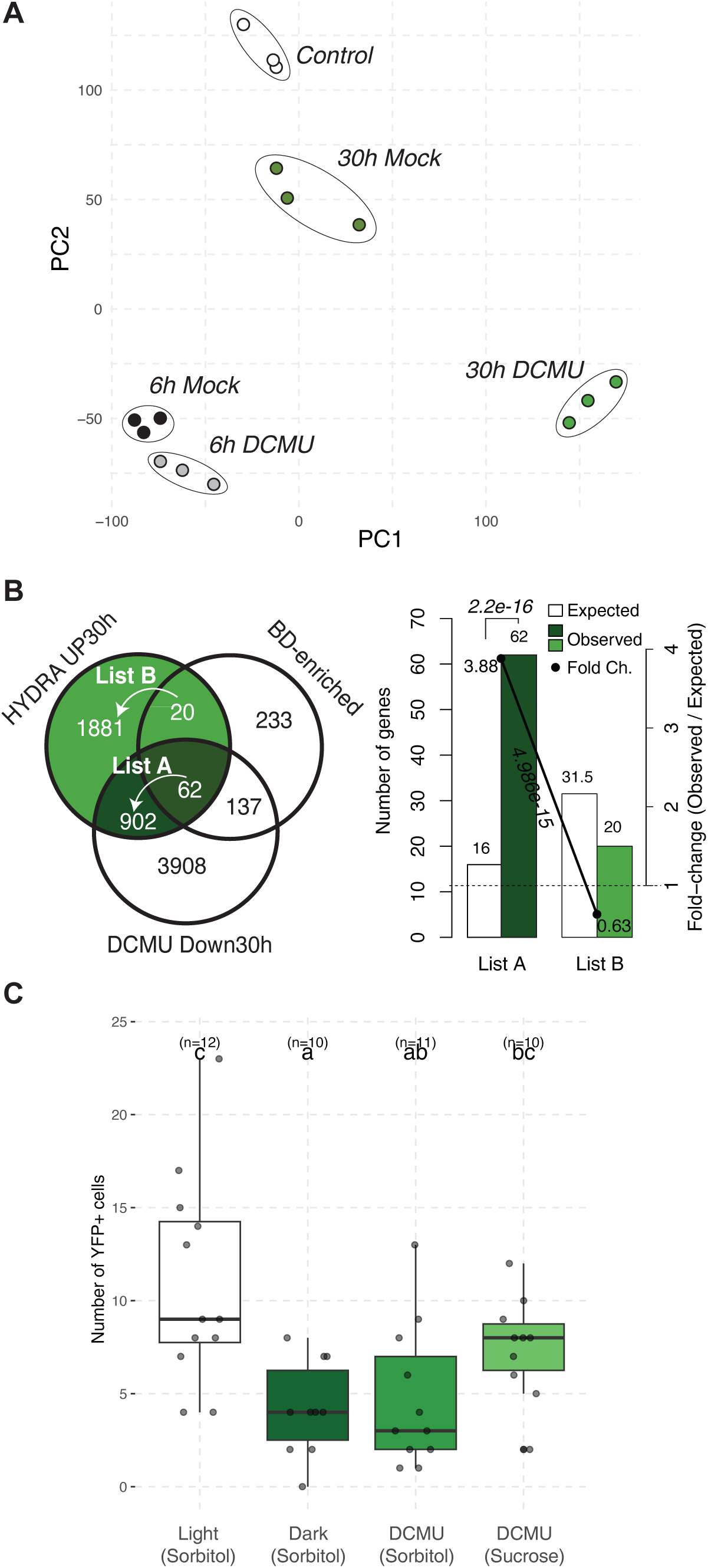
Transcriptome quality control and additional analyses (Related to Figure 4). (A) Principal component (PC) analysis of HYDRA RNA-seq samples. (B) Enrichment analysis comparing boundary-domain-enriched genes (BD-enriched, (Tian et al., 2014) with our datasets. (C) CYCB1;2-YFP signal in HYDRA explants at given conditions. Boxplots represent the distribution of the data, the center line indicates the median, the box corresponds to the interquartile range (25th to 75th percentile), and whiskers extend to 1.5 × the interquartile range. Different letters indicate statistical significance (P < 0.05). Sample size (n) is indicated.

**Supplementary Figure 6.**
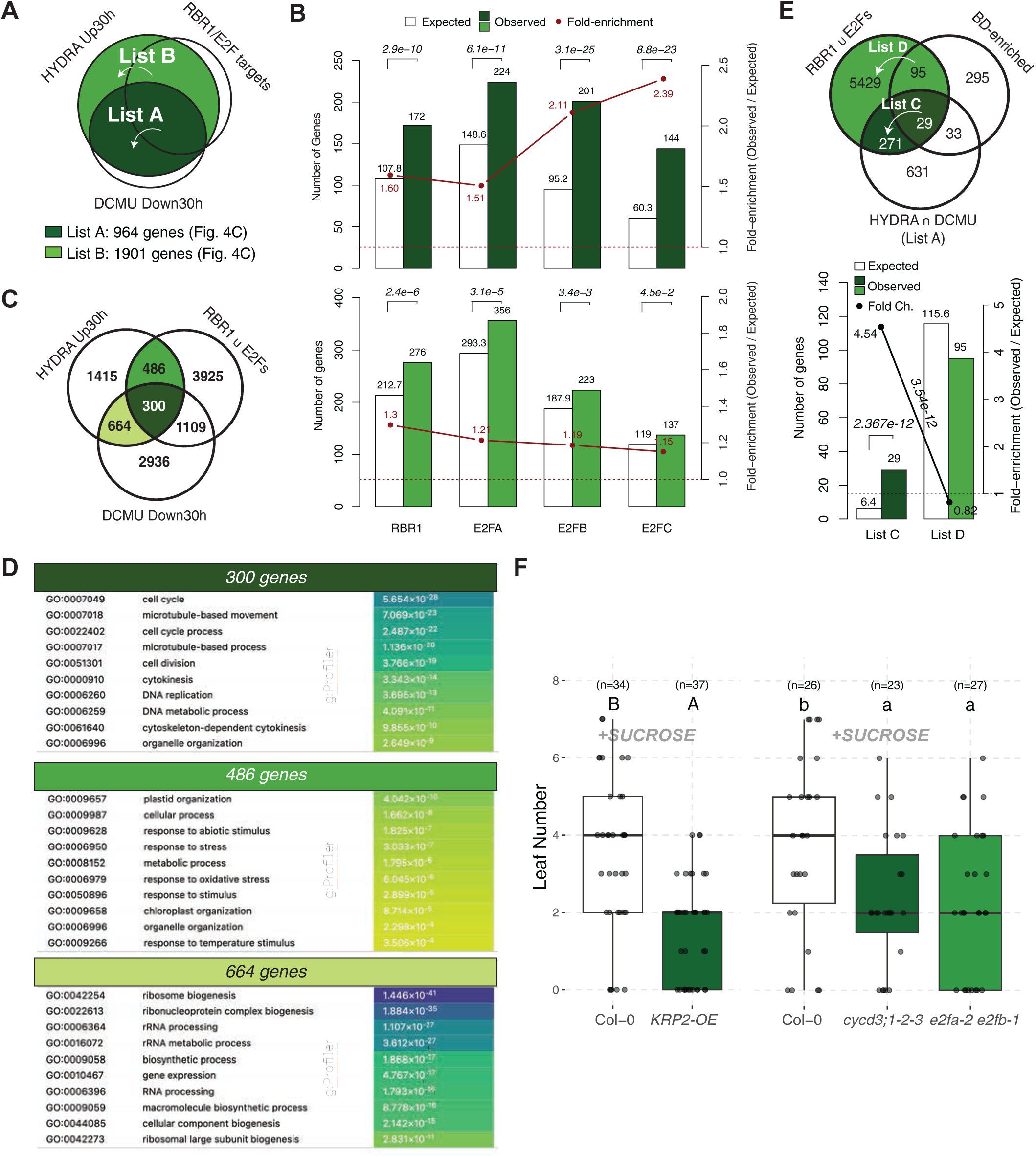
Comparison of HYDRA transcriptome with RBR1 pathway ChIP-seq datasets (Related to Figure 5). (A) Venn diagram comparing HYDRA dataset with RBR1/E2F-targets (Gombos et al., 2023). (B) Enrichment analyses comparing HYDRA dataset with RBR1/E2F-targets (Gombos et al., 2023). P values are indicated. (C) Venn diagram of genes listed from comparison of HYDRA dataset with combined RBR1 and/or E2Fs binding targets. (D) GO enrichment analysis of genes highlighted in (C) showing top 10 terms (ordered by FDR). (E) Venn diagram (top) and enrichment analysis (bottom) from comparison of HYDRA dataset with combined RBR1 and/or E2Fs binding targets and boundary domain (BD)-enriched genes (Tian et al., 2014). P values are indicated. (F) Shoot regeneration in cell cycle mutants on exogenous sucrose supplemented MS media. Boxplots represent the distribution of the data, the center line indicates the median, the box corresponds to the interquartile range (25th to 75th percentile), and whiskers extend to 1.5 × the interquartile range. Different letters indicate statistical significance (P < 0.05). Sample size (n) is indicated.

**Supplementary Figure 7.**
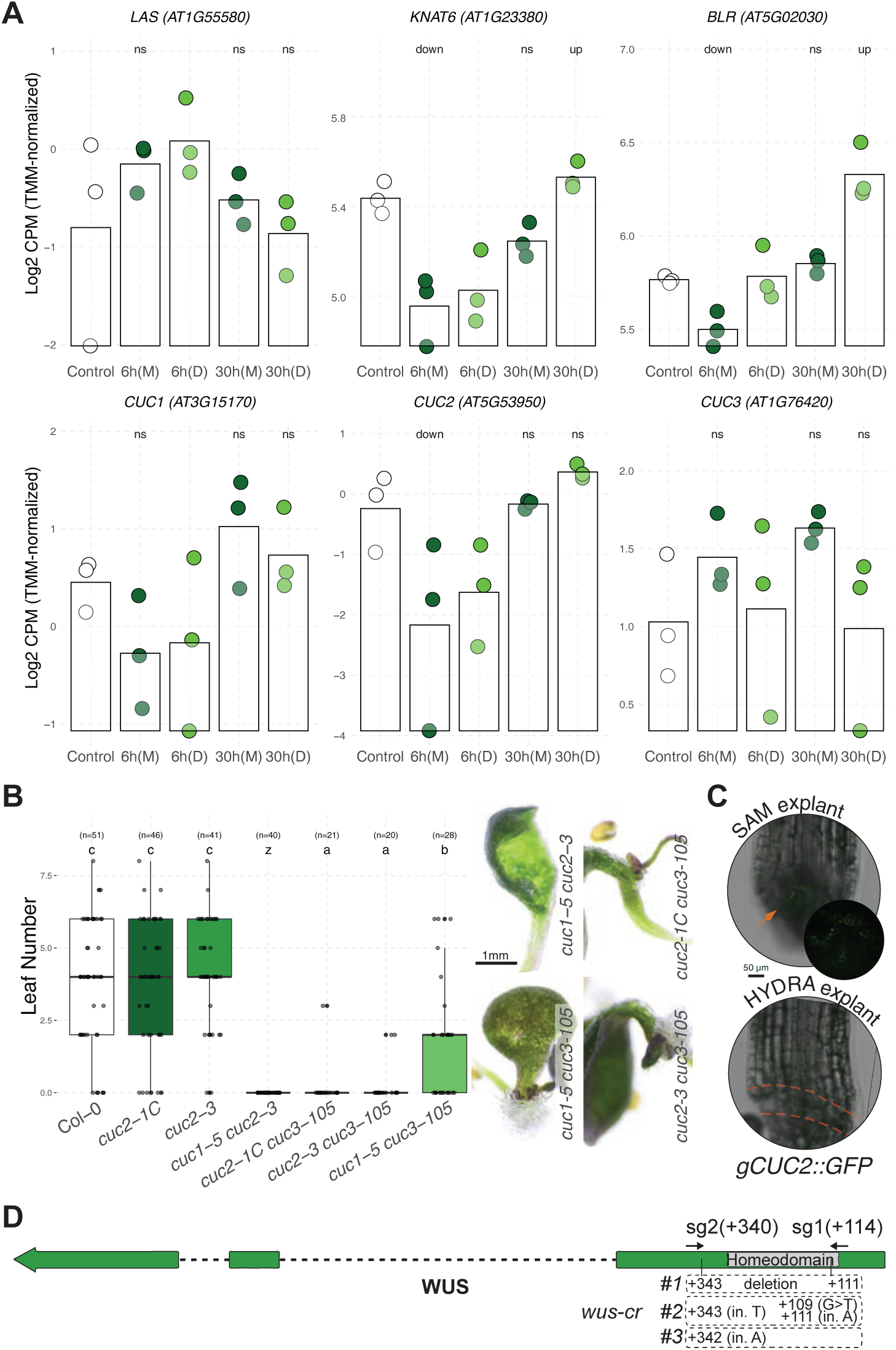
Meristem regulatory gene network in HYDRA (Related to Figure 6). (A) Expression of boundary-related genes *LAS*, *KNAT6*, and *BLR* and their upstream regulators, *CUC* family members in HYDRA. (ns: not significant, up: upregulated, down: downregulated, FDR<0.05). (B) Quantification (left) and representative images (right) of regeneration in *cuc* mutant combinations. (C) gCUC2::GFP signal immediately after HYDRA dissection. Images are shown as single optical section (top) and maximum projection (bottom). The region between orange dashed lines marks the boundary domain. (D) Genomic structure of *WUS* showing sgRNA target sites and sequencing confirmation of induced mutations. Dashed lines indicate introns, boxes exons, and arrowheads PAM sequences. Locations (bp) are given relative to start codon ATG. (in. insertion, > substitution). Boxplots represent the distribution of the data, the center line indicates the median, the box corresponds to the interquartile range (25th to 75th percentile), and whiskers extend to 1.5 × the interquartile range. Different letters indicate statistical significance (P < 0.05). Sample size (n) is indicated.

**Supplementary Figure 8.**
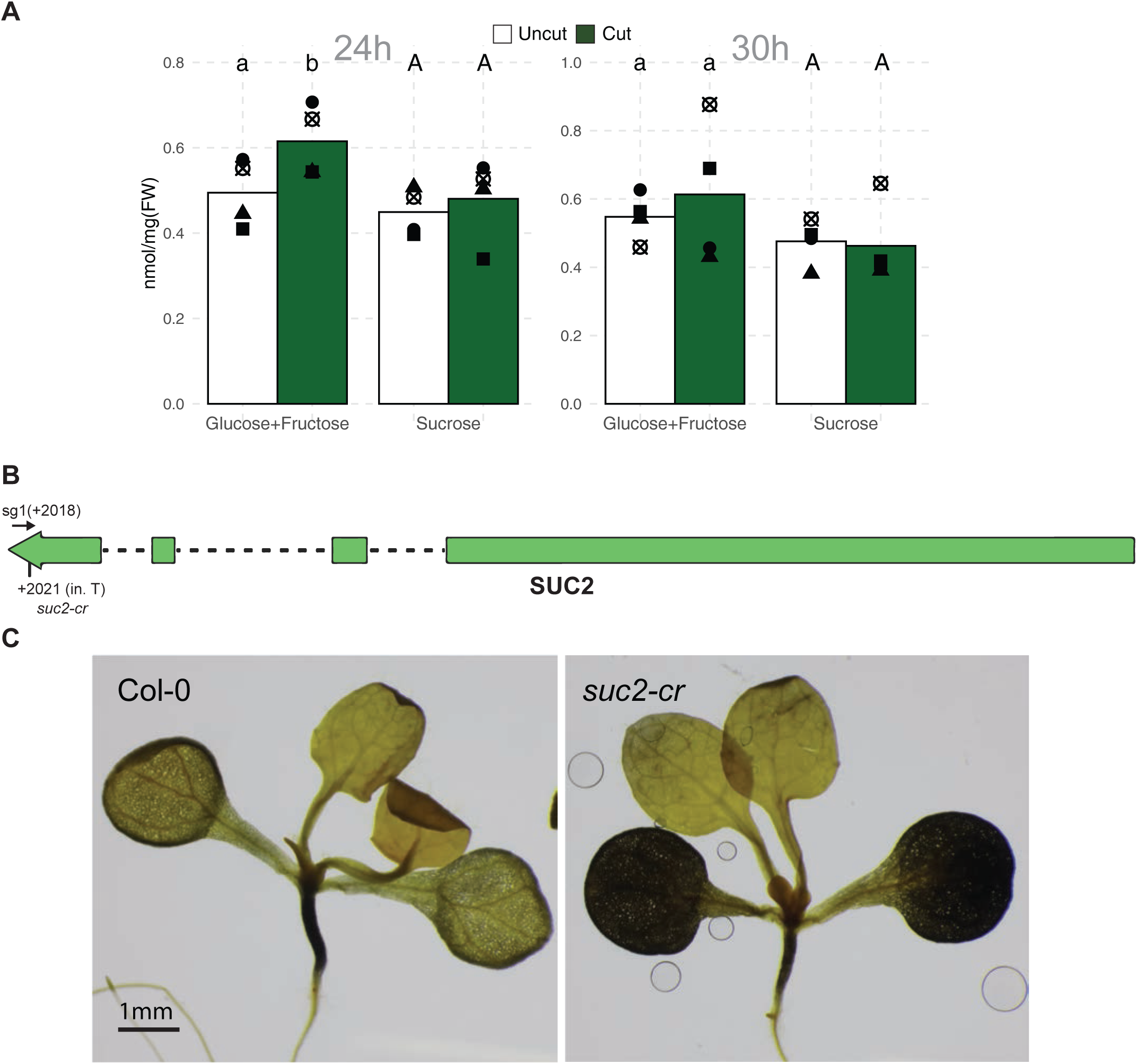
Sugar measurements in *B. rapa* and validation of *suc2-cr* mutant (Related to Figure 7). (A) (A) Sugar quantification in *Brassica rapa* boundary-enriched samples after dissection in comparison to control (uncut). Each dot corresponds to a biological replicate. Bars show average of these values. Different letters indicate statistical significance (P < 0.05). (B) Genomic structure of *SUC2* showing sgRNA target site and sequencing confirmation of induced mutation. Dashed lines indicate introns, boxes exons, and arrowhead PAM sequence. Locations (bp) are given relative to start codon ATG. (in. insertion) (C) Iodine staining of Col-0 and *suc2-cr* seedlings.

**Supplementary Table 1**. Gene sets used in this study.

HYDRA upregulated genes at 30h (Cut vs. Control, FC>1, FDR<0.05), given as carbon-dependent (List A) and carbon-independent (List B) (see Fig. 4C).

Boundary Domain Enriched Genes (expressed in our dataset) - (Tian et al., 2014)

E2FA bound (expressed in our dataset) - (Gombos et al., 2023)

E2FB bound (expressed in our dataset) - (Gombos et al., 2023)

E2FC bound (expressed in our dataset) - (Gombos et al., 2023)

RBR1 bound (expressed in our dataset) - (Gombos et al., 2023)

**Supplementary Table 2.** Gene Ontology Term Enrichment Results (FDR<0.05) for genes in List A (see Fig. 4C) (HYDRA up- DCMU-downregulated at 30h).

**Supplementary Table 3.** Plant material used in this study.

**Supplementary Table 4**. Chemical material used in this study.

**Supplementary Table 5.** Oligonucleotides used in this study.

**Supplementary Table 6.** sgRNAs used in this study.

